# Accurate isoform quantification in bulk and single-cell RNA-seq by integrating multi-platform sequencing data and sequence-to-expression models

**DOI:** 10.1101/2024.07.11.603067

**Authors:** Michael Apostolides, Jichen Wang, Ali Saberi, Benedict Choi, Tabea Mittmann, Ali Poursina, Arsham Mikaeili Namini, Darian Salehi, Gabrielle Perron, Albertas Navickas, Larisa M. Soto, Aiden Sababi, Hani Goodarzi, Hamed S. Najafabadi

## Abstract

Accurate quantification of transcript isoforms is crucial for understanding gene regulation, functional diversity, and cellular behavior. Existing RNA sequencing methods have significant limitations: short-read (SR) sequencing provides high depth but struggles with isoform deconvolution, especially in single-cell data with strong positional biases, whereas long-read (LR) sequencing offers isoform resolution at the cost of lower depth and platform-specific biases. Here, we introduce Multi-Platform Aggregation and Quantification of Transcripts (MPAQT), a probabilistic generative framework that integrates sequencing data across platforms while explicitly modeling their distinct data-generating processes and technical biases. Across extensive simulations and experimental benchmarks, MPAQT provides state-of-the-art transcript quantification from short-read data alone and further improves accuracy by jointly analyzing short- and long-read sequencing data in both bulk and single-cell transcriptomes. We show that long-read sequencing is particularly informative for resolving isoforms of genes with complex transcript structures, while integration with short reads mitigates biases and transcript dropouts inherent to long-read data. MPAQT can additionally incorporate sequence-to-expression models as informative priors that are iteratively refined by sequencing observations. Using models of transcript abundance based on RNA-binding protein recognition sequences and transcript co-expression patterns, we substantially improve isoform quantification from single-cell data and reveal a strong contribution of isoform-specific sequence features in untranslated regions to isoform abundance and exon usage. Together, our results show that complementary sequencing technologies and predictive models of transcript regulation jointly constrain latent transcript abundances, enabling accurate isoform-resolved quantification across bulk and single-cell RNA sequencing.

## Introduction

Nearly all protein-coding genes encode multiple transcript isoforms, resulting from a wide array of alternative transcription start sites (TSSs), transcription termination sites (TTSs), and/or alternative splicing of exons^1^. This isoform variation is a significant source of molecular and functional diversity, as proteins produced from different isoforms of the same gene can have distinct (and even opposite^2^) functions. Even transcript isoforms that encode the same protein can differentially affect cellular functions due to variations in mRNA localization, stability, and/or translation^3–5^. Thus, isoform abundances can reflect the biological state better than gene-level information.

Given the sequence similarity of transcript isoforms, accurate quantification of isoform abundances by RNA sequencing remains a major challenge. Isoform abundance quantification is fundamentally an inference problem with transcript abundances as latent variables that cannot be directly observed but must instead be reconstructed from incomplete and noisy sequencing observations^6^. Different sequencing technologies provide complementary views of the same underlying transcriptome, each with distinct strengths, biases, and ambiguities. Methods based on short-read (SR) sequencing can generate many reads at increasingly low costs, providing high sequencing depth. However, most short reads cannot be confidently assigned to a single isoform^6^. In contrast, long-read (LR) sequencing generates reads that almost capture full-length transcripts, allowing unambiguous assignment of most reads to a specific isoform, but currently suffers from lower throughput and technology-specific biases including transcript length, nucleotide composition, and incomplete transcript coverage^7, 8^. While existing methods have largely focused on developing specialized statistical frameworks for individual sequencing technologies, few integrate short- and long-read data, which primarily focus on splice junction refinement^9^, transcriptome assembly^10, 11^, or alternative polyadenylation site identification^12^. Methods that attempt to integrate SR and LR transcript quantification information do so without considering the technical biases of each platform or the ability to integrate additional sources of information about transcript abundance^13^.

These challenges become particularly severe in single-cell RNA sequencing, with widely used droplet-based protocols capturing only transcript termini, introducing strong positional biases that remove most sequence information required to distinguish transcript isoforms^14^. Due to limited isoform information recovered by existing computational approaches from such biased data^15^, most high-throughput single-cell RNA-seq analyses are performed at the gene level. Recent studies show LR sequencing recovers isoform-resolved transcriptomes from single cells (reviewed in ref^16^), but LR approaches remain constrained by sequencing depth and cost^17^, and existing approaches do not integrate key complementary information from SR and LR technologies.

Furthermore, a crucial but overlooked notion is that sequencing observations are not the only source of information about transcript abundance. Transcript sequences encode regulatory elements that influence transcript stability, localization, and processing. Thus, beyond sequencing data themselves, transcript abundance is constrained by regulatory information encoded within transcript sequences, which can be leveraged to obtain more accurate quantifications (e.g., see ref^18^). Recent predictive sequence-to-expression models, ranging from those relying on engineered sequence features ^19^ to deep neural networks^20–22^ and foundation models trained on RNA sequences^23, 24^, show increasing ability to capture transcript-level regulatory information. Such models provide information complementary to sequencing observations and could serve as informative priors for transcript abundance estimation. However, no existing framework integrates predictive sequence models with heterogeneous sequencing measurements within a unified statistical model for transcript quantification.

Here we introduce Multi-Platform Aggregation and Quantification of Transcripts (MPAQT), a probabilistic generative framework for isoform-resolved transcript quantification that integrates heterogeneous sequencing measurements and predictive sequence-to-expression models. MPAQT explicitly models the distinct data-generating processes and technical biases of different sequencing platforms, improving quantification accuracy from short reads alone as well as enabling joint analysis of short- and long-read sequencing in both bulk and single-cell RNA-seq. Furthermore, it incorporates sequence-to-expression models as probabilistic priors that are refined by sequencing observations, allowing complementary experimental and sequence-derived information to jointly constrain latent transcript abundances. Using extensive simulations, experimental benchmarks, and matched bulk and single-cell short- and long-read datasets, we demonstrate state-of-the-art transcript quantification and use the resulting abundance estimates to uncover sequence determinants of isoform regulation.

## Results

### The MPAQT framework

At the core of MPAQT is a generative model that connects latent transcript abundances to the observed counts for “observation unit” (OUs) (**Figure 1a**). Here, an observation unit is any entity that we can directly quantify from RNA-seq reads, defined in a technology-dependent manner. In long-read (LR) data, we can unambiguously assign most reads to one transcript isoform and define each isoform as one OU, allowing one-to-one correspondence between transcript isoforms and OUs. The expected count of each OU scales linearly with the abundance of its corresponding transcript, with factors such as transcript length or GC content affecting the slope of this relationship in a sequencing technology- or protocol-dependent manner. The observed count is then modeled as a sample from a Poisson distribution whose mean is the expected count determined by the transcript abundance and transcript-level covariates.

**Figure 1.**
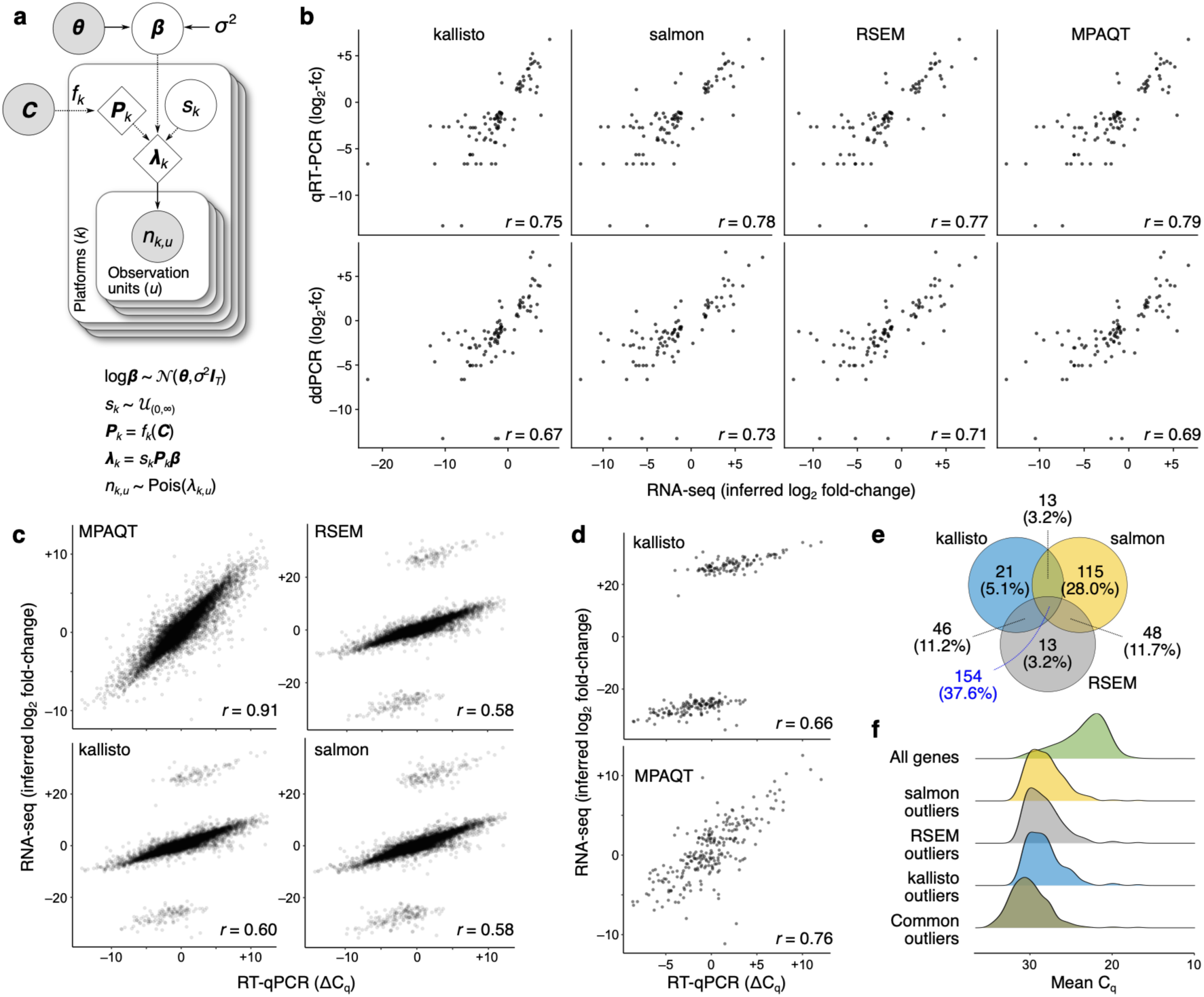
Overview of MPAQT and its performance for inference of gene-level abundances. **(a)** The generative model of MPAQT. Open circles, closed circles, and the diamonds represent latent variables, observed variables, and deterministic computations, respectively. ***β***: vector of transcript abundances; *sk*: library size for platform *k*; ***C***: set of transcript sequences; *nk*,*u*: number of reads mapping to the observation unit *u* in platform *k*. See **Methods** for description of other variables. (**b**) Comparison of inferred and ground truth measurements in the Quartet dataset for kallisto, salmon, RSEM, and MPAQT. Each dot represents inferred (x-axis) and ground truth (y-axis) log fold-change of one gene in a pair of cell lines. (**c**) Inferred log fold-change between MAQCA and MAQCB, calculated from TPM predictions by MPAQT, RSEM, kallisto, and salmon, plotted against the ground truth qPCR difference. Each point is one gene (*n*=14,956). **(d)** Top: kallisto’s outliers, isolated from other genes. Bottom: MPAQT’s inferences for kallisto’s outlier genes (*n*=234). **(e)** Venn diagram showing overlap of outliers among kallisto, salmon, and RSEM. **(f)** Density curves of mean of MAQCA and MAQCB Cq-values for each tool’s outliers and for all genes. For benchmarking results with the expanded filtering range, see **Supplementary Figure 2**. Data underlying this figure can be found in **Supplementary Data Table 1**.

Conversely, short-read (SR) RNA-seq reads map to multiple transcripts (or multiple genes), complicating the relationship between transcripts and OUs. Here, we use the equivalence classes (ECs)^6^ as the OUs. In this case, each transcript may be connected to multiple OUs, and each OU to multiple transcripts (extending the concept of ECs to long-read is also possible; see ref^11^). MPAQT models the expected count of each OU as a linear function of the abundances of all transcripts that may contribute to that OU (**Figure 1a**). The parameters of this function (i.e., the transcript-OU weights) are obtained through analysis of simulated short reads (see Methods for details), and can be adjusted to account for factors such as read positional biases.

By explicitly modeling the OU counts as probabilistic functions of transcript abundances, MPAQT provides a framework for joint analysis of data across multiple platforms that assay the same RNA sample, such as short- and long-read sequencing data (**Figure 1a**)—MPAQT infers transcript abundances by maximum a posteriori (MAP) estimation given a prior distribution for transcript abundances and the observed OU counts across all platforms. At the same time, MPAQT incorporates platform- and experiment-specific bias parameters that are directly learned from data, such as the length-, GC-, and positional biases.

### MPAQT enables state-of-the-art gene- and transcript-level quantification with short reads alone

To examine whether MPAQT might be broadly applicable to gene expression quantification, we began by benchmarking its performance against three leading SR analysis tools, salmon^12^, kallisto^6^, and RSEM^13^, for gene-level quantification using SR data alone. For this purpose, we started with a dataset from the Quartet Project, consisting of RNA reference materials from four cell lines from a monozygotic twin family for ratio-based transcriptomics benchmarking^25^. These materials exhibit small biological differences, allowing for fine-tuned assessment of RNA-seq-based measurements, with ratio-based gene-level ground truth obtained by RT-qPCR and ddPCR measurements for select genes, which was then compared to gene-level log fold-change of TPM (transcripts per million) calculated from SR RNA-seq data by different tools (gene-level TPM values in each sample were calculated by summing up transcript-level TPMs for each gene). Using RT-qPCR measurements as ground truth, MPAQT emerged as the best performing method; using ddPCR measurements as ground truth, MPAQT outperformed kallisto but not RSEM and Salmon (**Figure 1b**). However, given the relatively small number of genes present in this benchmarking dataset, the differences were not statistically significant.

For an expanded benchmarking, we used data from the MicroArray Quality Control (MAQC) project^14^, consisting of single-end RNA-seq data for two MAQC samples^15^, MAQCA (Universal Human Reference RNA, pool of 10 cell lines) and MAQCB (Human Brain Reference RNA), accompanied by RT-qPCR expression measurements for 18,080 protein-coding genes in the form of Cq-values. Of the genes that could be readily matched to quantifications from the RNA-seq-based methods based on their IDs, 14,956 genes had Cq-values between 11 and 32, a range deemed reliable in the original report^15^. For this subset, we observed excellent agreement between log fold-changes inferred by MPAQT and the ground truth differential expression (DE) values (Pearson r=0.91, **Figure 1c**). In contrast, for all other existing tools, we saw distinct outliers that were visibly separated from the remainder of the data points (**Figure 1c**). **Figure 1d** shows kallisto’s outliers, isolated from other, well-behaving genes. Interestingly, MPAQT preserves differential expression information for these outlier genes (**Figure 1d**). A similar trend is observed for outliers from salmon and RSEM (**Supplementary Figure 1**). These outliers represent ∼2-3% of genes in our data (**Figure 1e**) and correspond to genes with higher mean Cq-values (**Figure 1f**), suggesting they are genes with low expression; this trend toward higher mean Cq-values is even stronger for the 154 outliers shared among kallisto, salmon, and RSEM (**Figure 1f**), suggesting that low-abundance genes are commonly misquantified by the most commonly used existing tools. We also examined an extended filtering range (Cq-value between 8-35) to assess MPAQT’s performance on noisier qPCR measurements, allowing for an additional 1148 genes to be included in the analysis—surprisingly, MPAQT’s performance remained comparable to the more conservative filtering range (Cq-value between 11-32), whereas the number of outliers for the other SR tools approximately doubled (**Supplementary Figure 2**), highlighting the volatility of existing methods in the presence of very low-abundance genes.

To go beyond gene-level quantification and benchmark the performance of MPAQT in transcript-level quantification, we started with analysis of four simulated datasets: three with ground truth TPM values sampled randomly from an exponential distribution, and one with ground truth TPM values sampled from a distribution that was modeled after measurements from real RNA-seq data (see **Methods** for details). In all simulations, we observed that MPAQT substantially outperforms the other tools in terms of Pearson correlation and Root Mean Square Deviation (RMSD) (**Figure 2a-b**). The amount of variance in the ground truth log-TPMs that is captured by MPAQT (i.e., R^2^ between MPAQT inferences and ground truth) is ∼13%–32% higher than the next best method (13% for the data simulated for TPMs that are modeled after real RNA-seq measurements, and 32% for the data simulated for TPMs that are exponentially distributed).

**Figure 2.**
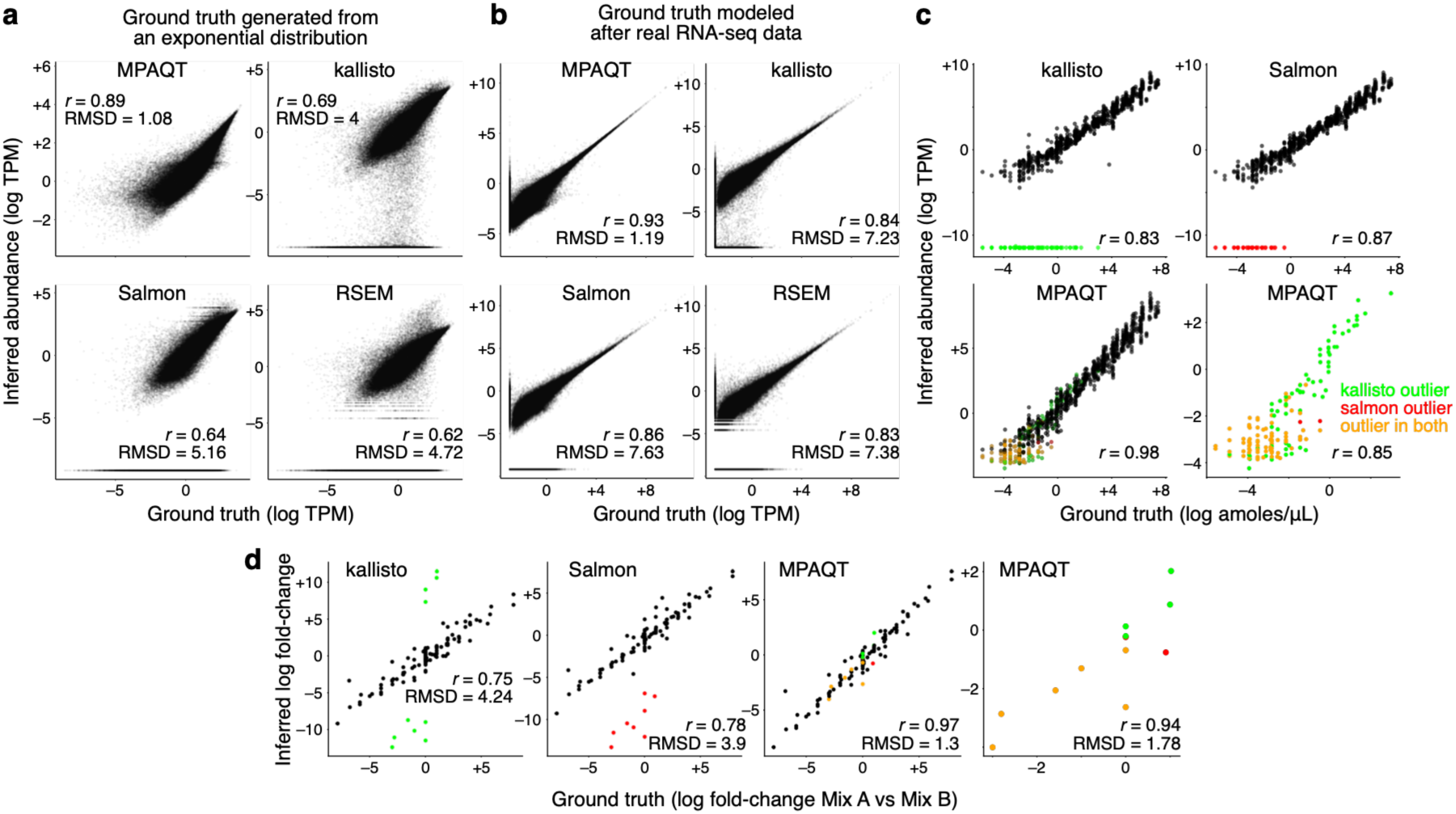
Accurate inference of transcript abundances from SR data using MPAQT. (**a**) Benchmarking of MPAQT and three other tools using simulated SR data with ground-truth TPMs generated from an exponential distribution. (**b**) Similar to (a), but for a simulated sample with ground-truth TPM values modeled after real data (see **Methods** for details). (**c**) Benchmarking results using Sequin transcripts with known concentrations. Each dot is one Sequin transcript in one cell line. Colored dots represent dropouts in kallisto and/or salmon inferences. The lower-right plot shows only the subset of transcripts that appear as dropout (TPM=0) by at least one tool. (**d**) Inferred log fold-change between two Sequin mixtures, plotted against ground truth log fold-change. Data underlying this figure can be found in **Supplementary Data Table 2**.

Interestingly, in these simulations, we observed many transcripts whose abundances were inferred as zero by all tools except MPAQT, despite their non-zero ground truth abundance. This observation can be replicated in real RNA-seq data: we benchmark MPAQT, kallisto, and Salmon on RNA-seq data from two cell lines, each in triplicate, spiked with Sequin transcripts with known abundances^26^. Using SR data alone, kallisto and Salmon’s estimates track linearly (in log-scale) with the true abundances for many transcripts (**Figure 2c**), but for a substantial portion, each report an abundance of zero—we refer to these cases as “dropouts”. Specifically, we observed dropout rates of 16.3% for kallisto (156 dropout events out of 960 measurements; 160 transcripts × 6 samples) and 9.2% for Salmon (88 droput events out of 960 measurements). In contrast, we did not observe any dropouts in MPAQT inferences, with MPAQT achieving markedly improved accuracy (Pearson correlation 0.98, compared to 0.83 and 0.87 for kallisto and Salmon, respectively). Importantly, MPAQT quantitatively recovers the ground truth abundances of kallisto and Salmon dropouts (**Figure 2c**, bottom-right).

This improvement in quantification can also be seen for transcript-level DE analysis. Specifically, we made use of the fact that the two cell lines in the above analysis were spiked with two different Sequin mixes which have different ground truth concentrations for each of the spike-in transcripts (Mix A for cell line H1975, and Mix B for cell line HCC827), and performed DE analysis between these two cell lines for the 160 Sequin transcripts. When compared to the ground truth log-FC of Sequins, both Salmon and kallisto showed distinct outliers for which the differential abundance was not correctly inferred (colored points in **Figure 2d**). In contrast, MPAQT did not show any outliers, and overall had substantially higher accuracy in inferring differential abundance (Pearson correlation 0.97, compared to 0.75 and 0.784 for kallisto and Salmon, respectively), including for the subset of Salmon and kallisto outliers (Pearson correlation 0.94, **Figure 2d**). Together, these analyses suggest that MPAQT can rescue genes and transcripts misquantified by commonly used SR analysis tools.

### Modeling read positional biases enables isoform-level quantification in short-read single-cell data

The flexibility of MPAQT’s generative framework allows it to infer from data potential sources of bias and incorporate them in its model. This ability is especially important in analysis of data from common single-cell RNA-sequencing technologies, in which extreme 5’ or 3’ positional biases render isoform abundance inference challenging. These biases modify the data generation process by affecting the transcript-OU conditional probabilities; for example, in 3’ single-cell data, OUs that represent regions closer to 3’ end of a transcript are more likely to be observed. Similarly, when the same OU is compatible with multiple transcripts, transcripts whose 3’ ends are overall closer to the region represented by that OU have higher likelihood of being the origin of the reads mapping to that OU (**Supplementary Figure 3**). Thus, positional biases violate the core assumption that reads are uniformly distributed along transcripts and, thus, transcript-OU weights are proportional to the represented sequence length. MPAQT learns these biases from the RNA-seq data (see **Methods** for details) and uses them to correct transcript-OU weights. **Figure 3a-b** show an example, wherein by learning the relationship between read position and capture probabilities, inferred isoform usage profile can be corrected (read positional biases are learned from pseudobulked reads across the entire scRNA-seq dataset, and then used for isoform abundance inference in predefined cell clusters; see **Methods** for details).

**Figure 3.**
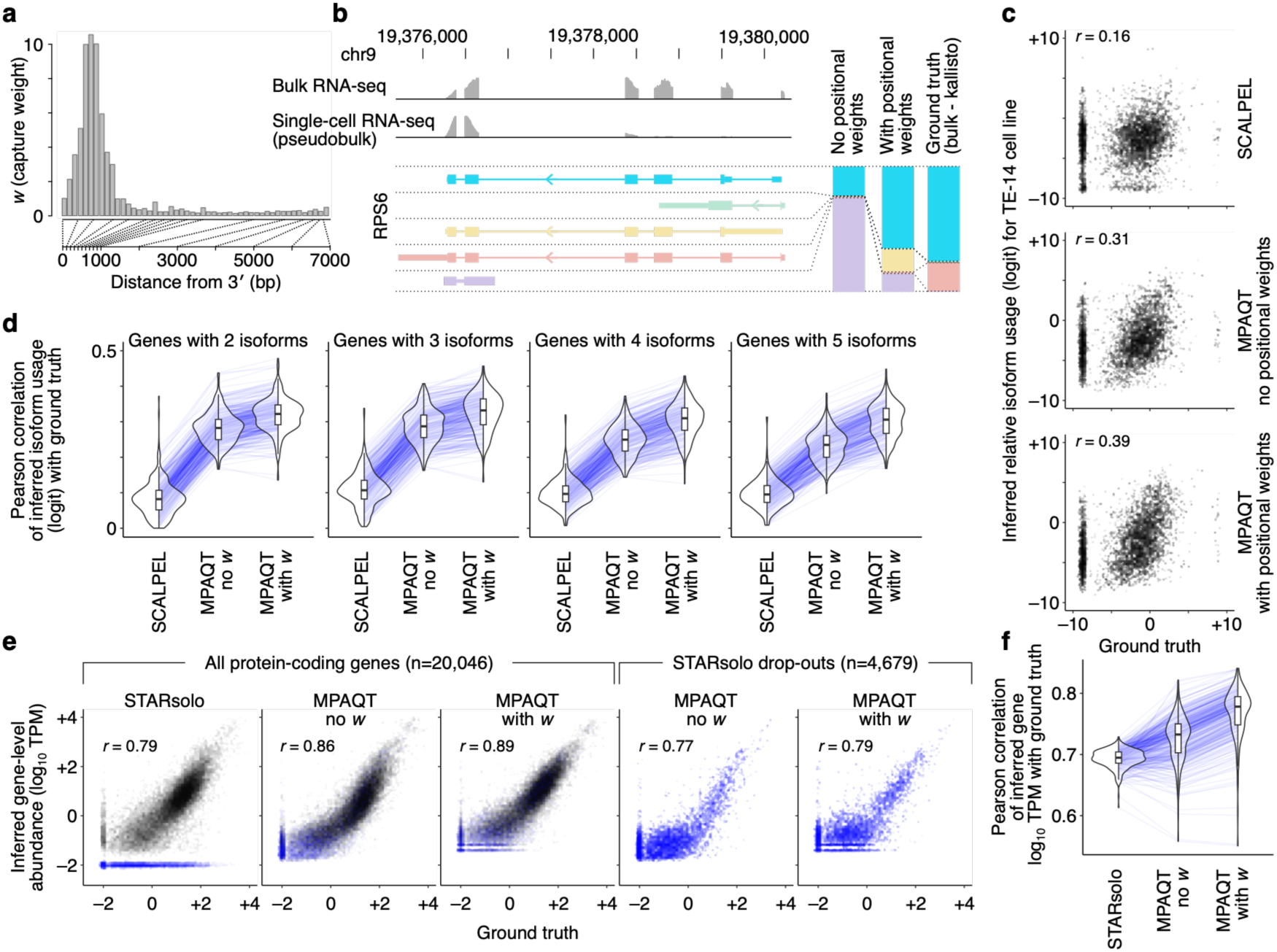
Isoform quantification in 3’-biased single-cell data with MPAQT. (**a**) Example positional bias profile inferred by MPAQT (based on single-cell data of CCLE cell line mixture^27^; SRR12557892). The x-axis shows the distance from 3’ end of the transcript (binned to represent equally distanced quantiles). (**b**) Bulk and single-cell read profiles for *RPS6* gene. *RPS6* isoforms are shown with colors matching the entries in the stacked bar plot on the right, which shows the relative usage of each isoform as inferred from single-cell or bulk data (the latter represents the ground truth). (**c**) Relative isoform usages inferred from pseudobulked single-cell data for TE-14 cell line (y-axis) vs ground truth values (x-axis, based on bulk RNA-seq). A pseudocount of 0.01 with a small jitter is added to all TPM values before relative isoform usage calculation to avoid undefined logits. Transcripts from protein-coding genes with 5 isoforms and gene-level TPM >1 are included in these example plots. TE-14 is chosen for this example as it constituted the largest number of cells in the CCLE mixture. (**d**) Systematic comparison of inferred vs ground truth relative isoform usages across 185 cancer cell lines (inferred from cell line-level pseudobulk of single-cell data). Each line represents one cell line. MPAQT no *w*: MPAQT without positional bias learning; MPAQT with *w*: MPAQT with positional bias learning. Results are shown separately for genes with varying isoform complexity (defined as the number of isoforms encoded by each gene). Only protein-coding genes with gene-level TPM >1 are included in the calculation of Pearson correlations. Violin plots represent the estimated distribution of Pearson correlation values for each inference tool. (**e**) Comparison of gene-level abundances inferred from single-cell data (pseudobulk) and ground truth (bulk RNA-seq) for TE-14 cell line. Blue dots represent genes with STARSolo-inferred TPM=0 (a pseudocount of 0.01 with a small jitter is added to all values for the purpose of plotting on log-scale). The MPAQT inferences for these STARSolo dropouts are separately shown in the plots on the right. (**f**) Systematic comparison of inferred vs ground truth gene-level abundances, similar to panel (d). Data underlying this figure can be found in **Supplementary Data Table 3**.

To systematically evaluate the effectiveness of MPAQT’s single-cell isoform abundance inference using SR data alone, we benchmarked MPAQT with and without positional bias correction using SR single-cell data^27^ generated from a mixture of 185 cell lines from the CCLE collection^28^ (pseudobulked at the level of cell line), with bulk RNA-seq of each individual cell line (quantified using kallisto) as ground truth. We observed that modeling read position biases almost invariably improves relative isoform quantification by MPAQT, as evident from comparison of inferred vs ground truth relative isoform usages (**Fig 3c-d**). MPAQT also significantly outperformed SCALPEL^15^, which, to our knowledge, is the only existing tool specifically designed for isoform quantification from 3’ short-read single-cell RNA-seq data (**Fig 3d**). Interestingly, modeling read positional biases also improves gene-level quantification: using kallisto bulk gene-level aggregation of isoform TPMs as ground truth, MPAQT’s aggregated gene abundance even outperforms STARsolo^29^ (**Fig 3e-f**), which consistently ranks among the most accurate existing single-cell quantification methods^30–32^. Notably, we observed a subset of genes with STARsolo-inferred TPM of 0 but a wide range of ground-truth TPMs (blue dots in **Fig 3e**), which parallels our findings with dropouts in bulk benchmarking experiments; MPAQT quantitatively recovers these genes with similar accuracy to all other genes. Together, these results highlight state-of-the-art performance of MPAQT in both isoform and gene quantification from SR single-cell data. In the next section, we will show how this accuracy further improves by combining SR and LR data.

### Joint analysis of short and long reads improves isoform-level quantification in bulk and single-cell data

MPAQT’s generative model enables joint analysis of multiple distinct sequencing assays, such as both SR and LR data, from the same RNA sample. To evaluate this ability, we started with analysis of Sequin transcripts^26^ as in the previous sections, but with addition of matching PacBio (PB) or Oxford Nanopore Technology (ONT) long-read data. As SR-only quantifications were already near-optimal in this dataset (Pearson correlation 0.98, **Fig 2c**), expectedly we did not observe any improvement by adding SR+LR data (**Fig 4a**). It is nevertheless noteworthy that this performance was substantially higher than that of miniQuant^13^, a recent method that uses a gene-specific weighted combination of SR-based and LR-based likelihoods to estimate isoform abundances in bulk data (Pearson correlation 0.87 with PB and 0.91 with ONT data for miniQuant, compared to 0.98 for MPAQT; **Fig 4a**). We saw similar results in another benchmarking dataset, the Universal Human Reference RNA (UHRR), which includes ground truth abundances for 906 transcripts based on TaqMan RT-qPCR measurements^33^. Specifically, MPAQT inferences from SR data alone achieved Pearson correlation of 0.79, which only improved to 0.80 with addition of PB or ONT long-reads (**Supplementary Fig 4**). This was still substantially more accurate than SR-only inferences from kallisto, Salmon, and RSEM (Pearson correlations between 0.71-0.72) and SR+LR inferences with miniQuant (Pearson correlation 0.73-0.74, **Supplementary Fig 4**). We reasoned that, while demonstrating the superiority of MPAQT, these datasets are not challenging enough to showcase the added benefit of including long reads, as SR data can be enough for accurate RNA abundance inference with MPAQT. Specifically, LR data are expected to be useful in more complex scenarios, e.g., to resolve read ambiguities in (a) genes with complex splicing landscapes, or (b) single-cell data where short reads mostly come from 3’ or 5’ ends, inadequate to resolve isoforms even when accounting for positional biases.

**Figure 4.**
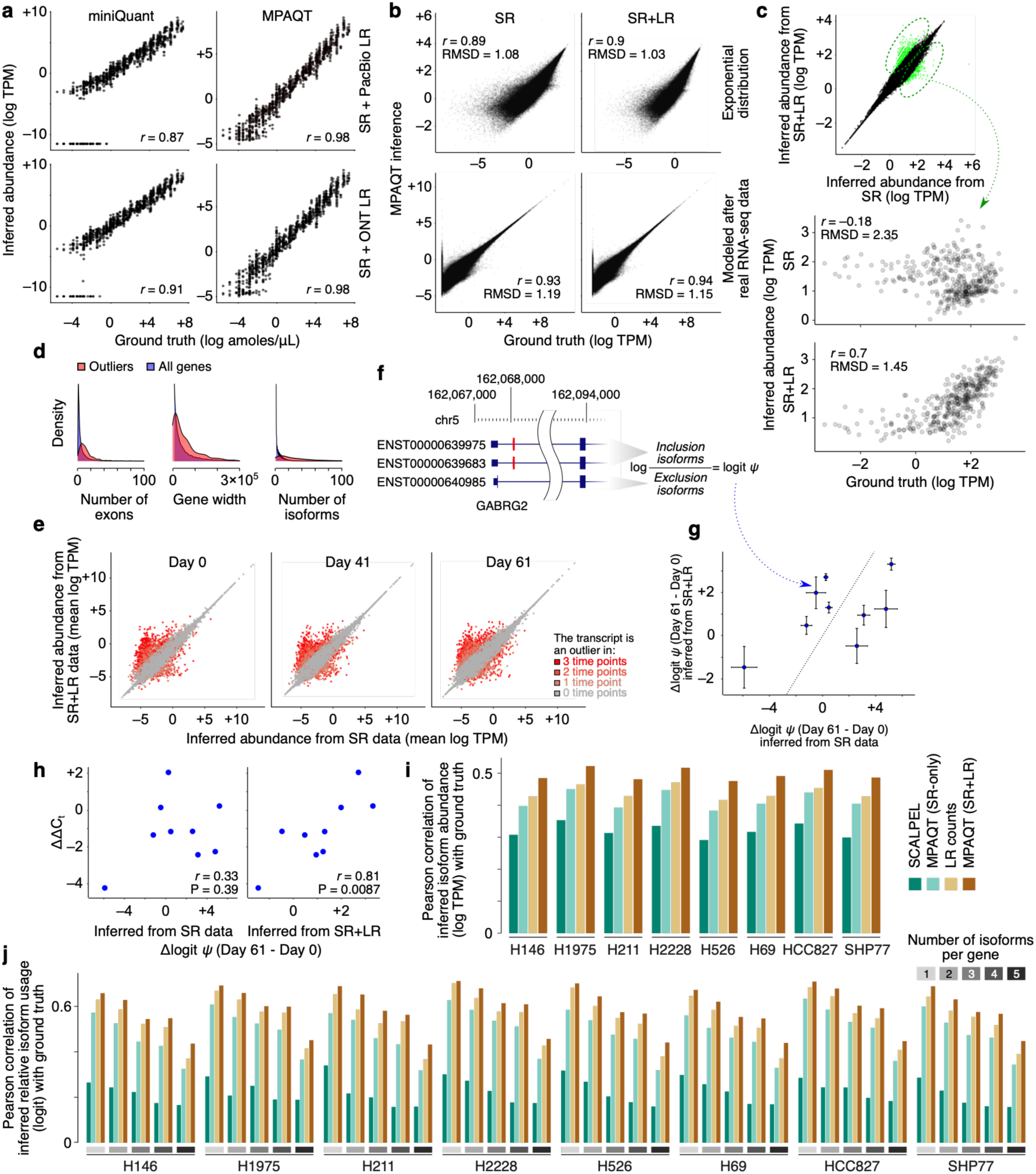
Isoform abundance inference from SR-LR joint analysis in bulk and single-cell data with MPAQT. (**a**) Inferred abundances, using either SR + PacBio LR data (top row) or SR + ONT LR data (bottom row), plotted against known concentration for Sequin transcripts, both for miniQuant (left) and MPAQT (right). (**b**) Performance of MPAQT using SR data alone (left) or SR+LR data (right), on simulated data generated from ground truth TPMs with an exponential distribution (top) or modeled after real RNA-seq data (bottom). SR plots are identical to the top-left plots in Figure 2a-b, and are repeated here for easier comparison to SR+LR. (**c**) Top: Comparison of SR vs. SR+LR inferences for one dataset simulated from an exponential distribution (see Supplementary Figure 3 for more simulation repeats). Transcripts with substantially different inferences are highlighted (outlier analysis based on Mahalanobis distance >6.36, equivalent to upper-tail P<10^-10^ for normally distributed data). Bottom: Comparison of inferred vs. ground truth TPMs for transcripts that are significantly differentially quantified (i.e., transcripts highlighted in the top panel). (**d**) Distributions of number of exons, gene width, and number of isoforms for genes encoding the transcripts that are differentially quantified between SR and SR+LR analyses. The distributions for all genes are also shown, for comparison. (**e**) Comparison of inferred TPMs based on SR data alone (x-axis) vs. SR+LR data (y-axis) in each of the three time points during neuronal differentiation. For each measurement, the mean of two replicates is used. Each data point is one transcript, with the dot color representing the number of time points in which the inferred abundance of the transcript differs significantly between SR and SR+LR measurements (Mahalanobis distance >6.36). (**f**) Quantification of cassette exon percent-spliced-in (PSI) from transcript isoform abundances. An example cassette exon for gene GABRG2 is shown for demonstration (shown in red). PSI (shown with *ψ*) is calculated as the sum of abundances of the isoforms that include the exon divided by that of all isoforms. (**g**) Top cassette exons for which the inferred change in PSI between days 0 and 61 differ significantly depending on whether SR or SR+LR quantifications are used. (**h**) Scatterplot of qPCR-based differential PSI quantification (y-axis) vs. differential PSI values inferred from SR data alone (x-axis of the left plot) or SR+LR data (x-axis of the right plot). (**i**) Benchmarking absolute isoform abundance quantification (log-TPM) from single-cell data (pseudobulked) in the LongBench dataset. Results are shown separately for each cell line in the LongBench cell line mixture. (**j**) Similar to (i), but for relative isoform usage quantification. Results are stratified by the number of isoforms per gene. Data underlying this figure can be found in **Supplementary Data Table 4**.

To examine the first scenario (i.e., genes with complex splicing landscapes), we expanded our simulated SR bulk datasets by simulating LR counts using the same ground truth TPMs. Specifically, we generated the LR full-length transcript counts by sampling from independent Poisson distributions, with each transcript’s ground truth TPM used as its Poisson distribution mean (adjusted to obtain ∼200K transcript counts per sample). Then, the combination of simulated SR data and LR counts was used as input to MPAQT, and the results were compared between SR-alone and SR+LR quantifications (**Figure 4b**). When MPAQT’s inferences from SR data are directly compared to those from SR+LR data, we can identify a subset of transcripts whose quantified abundances differ substantially between the two measurements (**Figure 4c**), despite the modest depth of the simulated LR datasets. For this subset, we see substantial improvement in ground truth agreement for SR+LR data in terms of agreement with ground truth (Pearson correlation 0.67 to 0.71 for SR+LR, –0.26 to –0.18 for SR, **Figure 4c** and **Supplementary Figure 5a**), suggesting LR data can substantially improve transcript quantification when combined with SR data.

Interestingly, although the simulation replicates were based on independent (uncorrelated) ground truths, among the 490 transcripts for which inclusion of LR data had a significant effect in at least one simulation, ∼30% were identified in more than one simulation (**Supplementary Figure 5b**), suggesting that these transcripts may have intrinsic features that make them sensitive to the presence/lack of LR data. We find that the genes encoding these transcripts have more exons, are longer, and have more isoforms compared to other genes (**Figure 4d**). Furthermore, pathway enrichment analysis revealed a significant and recurrent enrichment of “nervous system development” among the LR-sensitive genes in all replicates (FDR<7×10^-5^ for the three replicates, based on g:Profiler^16^). This finding is consistent with previous reports showing that genes preferentially expressed in the nervous system tend to be longer, have more exons, and exhibit more complex splicing patterns compared to other tissues ^4, 17^, and suggests that joint analysis of SR and LR data using MPAQT is particularly beneficial to the quantification of isoforms involved in nervous system development. To test this hypothesis, we analyzed human embryonic stem cells (hESCs) undergoing *in vitro* differentiation toward cortical neurons (**Supplementary Figure 6**), by joint SR and LR RNA-seq (PB) from cells collected at days 0, 41, and 61 since the start of growth in neural induction medium (see **Methods** for details). We found 6309 transcripts whose inferred abundances based on SR+LR analysis deviated substantially from abundances inferred from SR-only analysis in at least one of the three time points (Mahalanobis distance >2.32, equivalent to upper-tail P<0.01 for normally distributed data). Even at a substantially stricter cutoff (Mahalanobis distance >6.36, equivalent to upper-tail P<10^-10^), there were still 2459 transcripts whose quantifications differed significantly between SR+LR and SR-only in at least one time point (**Figure 4e**). In line with findings from our simulations, these transcripts corresponded to longer genes with more exons and more isoforms (**Supplementary Figure 7**).

To validate the higher accuracy of SR+LR transcript quantifications, we first used the inferred transcript abundances to estimate the percent-spliced-in (PSI) of alternatively spliced exons (**Fig 4f**), then selected nine cassette exons whose day-0-to-day-61 change in PSI differed significantly between SR and SR+LR measurements. (**Figure 4g**)—more accurate transcript quantifications are expected to result in more accurate PSI quantifications, of which the latter can be validated by RT-qPCR using junction-specific primer pairs. The selected cassette exons were mostly from neuron-specific genes and often associated with nervous system and/or psychiatric disorders (**Supplementary Figure 8**). For these exons, we quantified the true change in PSI using RT-qPCR and compared to differential PSI measurements from SR or SR+LR analyses. As shown in **Figure 4h**, we found that SR+LR analysis provides substantially more accurate estimates of differential PSI (*r=*0.81, P=0.0087) compared to SR-only analysis (*r=*0.33, P=0.39). These results further support joint analysis of SR+LR data as crucial for accurate estimation of transcript abundance, especially for transcripts involved in neuronal function and related diseases.

To examine the second scenario (i.e., whether LR sequencing improves transcript quantification in single cell data), we examined the MPAQT’s performance on joint SR+LR single-cell data. We used the recent LongBench reference dataset^34^, which contains SR and LR single-cell data for a mixture of eight cancer cell lines and matching bulk data for each individual cell line as ground truth. We observed consistent improvement in pseudobulk isoform abundance inference when single-cell SR and LR data were jointly analyzed: in all eight cell lines, pseudobulked single-cell SR+LR (with positional bias correction) significantly outperformed both SR alone and LR alone (mean Pearson correlation with ground truth: 0.43 for SR with positional bias correction, 0.47 for LR, and 0.50 for LR+SR with positional bias correction; **Fig 4i**). Similar improvements were observed for relative isoform usage inference across genes with varying isoform complexity (**Fig 4j**). Together, these results underline joint SR+LR analysis as the optimal strategy for isoform quantification.

### Integration with short reads mitigates biases inherent to long-read data

In our analysis of SR and PacBio LR data from differentiating neurons, we noticed an overall moderate correlation between SR-only TPM quantifications and LR raw counts (Pearson correlation of 0.3 for log-scaled values, **Fig 5a**), although MPAQT’s SR-based isoform quantifications correlated more strongly with LR raw counts than did other SR-based quantification tools (**Fig 5b**). We confirmed that SR-only quantification inaccuracy was not the cause of this moderate correlation between SR-only and LR-only quantifications: MPAQT’s SR-only TPM quantifications for synthetic mRNAs that were spiked in our samples correlate highly with ground truth concentrations (Pearson correlation 0.96; **Fig 5c**), while LR counts were only moderately correlated with ground truth (Pearson correlation 0.58; **Fig 5d**). Other PB and ONT datasets also exhibited this same moderate correlation with ground truth (**Supplementary Fig 4**).

**Figure 5.**
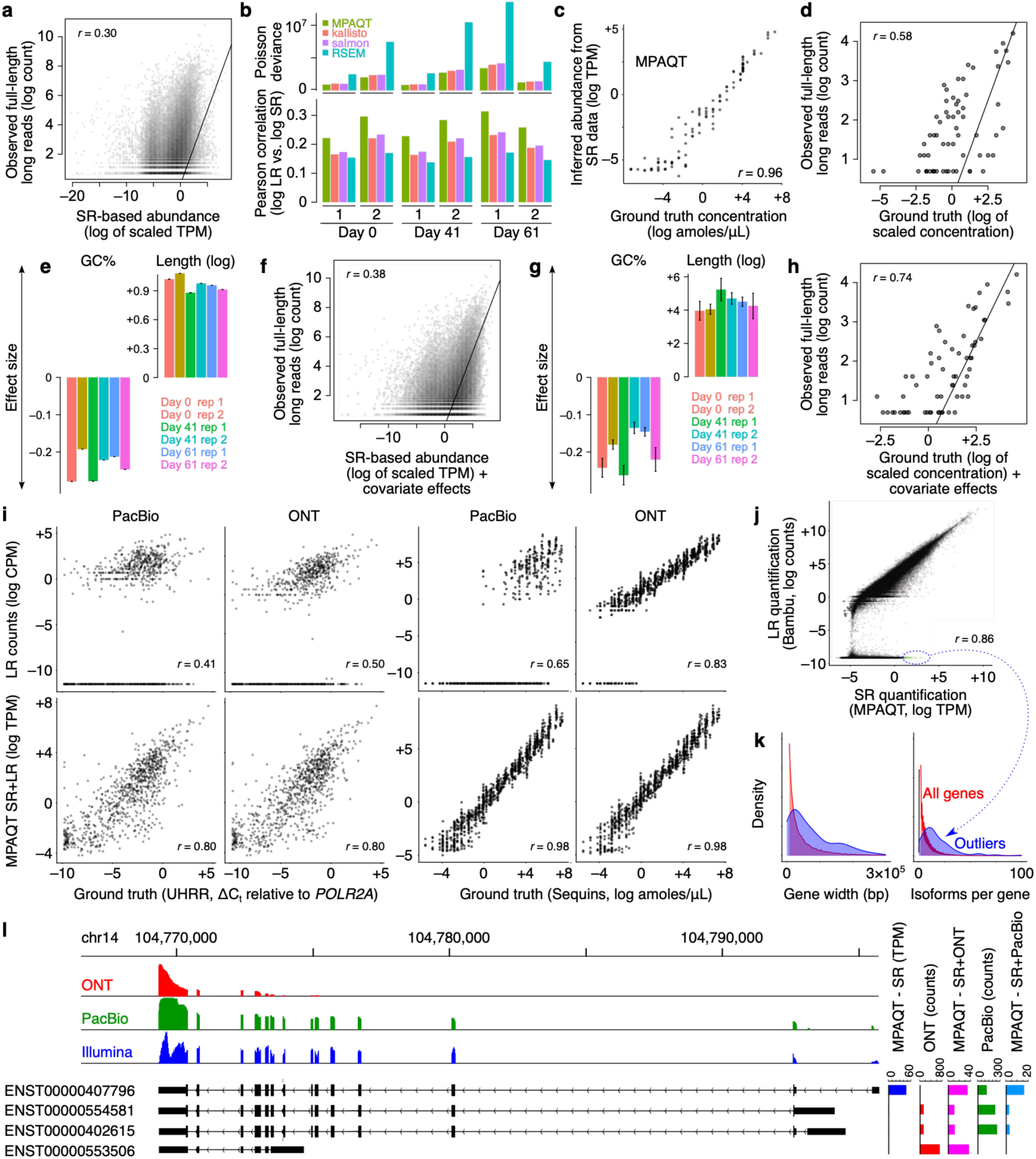
LR-only quantifications are affected by sequence-dependent biases and dropouts. (**a**) Full-length LR counts vs MPAQT SR abundance estimates for endogenous isoforms in the neuronal differentiation dataset. Each point represents one transcript in one sample (six samples combined in one plot). The SR-based abundances were scaled separately for each sample to match the scale of LR counts. Data points with LR count of zero were removed to allow log-scale plotting of the y-axis. (**b**) Comparison of SR-based inferences of different tools and full-length LR counts for endogenous transcripts. Left: Pearson correlation between LR log-counts and SR-based inferences, for each tool and each time point/replicate separately. Right: Poisson deviance is shown as an alternative measure of goodness of fit. (**c**) MPAQT SR-based estimates vs ground truth concentration for 107 spike-in transcripts. Each point represents one spike-in RNA in one sample/replicate. (**d**) Same as (c), but with full-length LR counts instead of SR-based estimates. Ground truth concentrations were scaled to match the scale of LR counts in each sample before pooling the data points together in this graph. (**e**) The effect of GC content and length on LR counts of endogenous transcripts, estimated by including them as covariates in a Poisson regression with log-scale SR-based inferences as the regressor. (f) Similar to (a), but the SR-based abundances are adjusted by adding the effect of covariates (transcript GC content and length). (g) Similar to (e) but showing the coefficients of GC content and length based on spiked-in transcripts, with log-scale ground-truth RNA concentration as the regressor. (**h**) Similar to (d), but the ground truth concentrations are adjusted by adding the effect of covariates (transcript GC content and length). (**i**) Comparison of LR counts (top row) or MPAQT SR+LR inferences (bottom row) vs ground truth in the UHRR (left) or the Sequin (right) datasets. For each dataset, PacBio and ONT LR counts are analyzed and shown separately. (**j**) Comparison of ONT LR counts (y-axis) and MPAQT SR quantifications (x-axis) for endogenous transcripts in HCC827 cells (data from ref^26^). Isoforms that, based on SR data, have high TPM values but are assigned zero long reads are highlighted by the oval (outliers). (**k**) Comparison of gene width and isoform counts between outlier genes (with LR dropout) and all genes. (**l**) LR and SR profiles of *AKT1* gene. *AKT1* isoform structures are shown at the bottom. The inferred abundances of each isoform based on different approaches are shown on the right. Data underlying this figure can be found in **Supplementary Data Table 5**.

In our neuronal differentiation dataset, deviation between LR and SR sequencing data for endogenous (genomic) isoforms can be at least partially explained by biases in the LR sequencing technology itself: longer and GC-poor transcripts are more likely to be captured in our PB LR data (**Figure 5e**). These biases also exist in other datasets we analyzed, including both PB and ONT data, but can be experiment- or platform-specific, as effect sizes differ between experiments (**Supplementary Fig 9**). Once we account for the biases introduced by these factors, we see a better agreement between SR and LR data (Poisson likelihood ratio test P<10^-16^, **Figure 5f**). Importantly, we observe these same biases when comparing LR spike-in counts to their ground truth concentrations (**Figure 5g-h**). MPAQT’s statistical model enables the inference of sample-specific sources of bias and incorporates them in its framework for integration of LR and SR data (SR+LR analyses presented in previous sections explicitly model GC and length biases; see **Methods** for details).

In addition to GC and length biases, we also observed frequent “dropouts” in LR data of spike-in RNAs, i.e., transcripts that, despite relatively high ground truth abundance, were assigned no long reads. This dropout issue was almost invariably resolved by joint analysis of SR and LR data with MPAQT, as shown in **Fig 5i** (but not with miniQuant, **Supplementary Fig 4**). These dropouts are also seen when comparing LR-only and SR+LR inferences for endogenous transcripts, where many transcripts with no assigned long reads receive large TPM values when SR data are incorporated (**Fig 5j** shows this effect in ONT data from HCC827 cell line). Interestingly, these LR-specific dropouts are enriched for broader gene widths and more annotated isoforms (**Fig 5k**), suggesting that SR data provide additional information about the isoform usage of complex genes. Anecdotally, we observed that this issue may stem from mis-assignment of long reads. **Fig 5l** shows AKT1, a critical component of the PI3K/AKT signalling pathway, as an example, where ONT long reads identified the short isoform ENST00000553506.5 as the most abundant isoform, whereas both Illumina short reads and PacBio long reads supported longer isoforms. ONT reads appear to have a 3’ coverage bias, potentially due to read truncation, explaining the spurious quantification of truncated isoforms (**Figure 5I**)—similar biases can be observed in other genes (**Supplementary Figure 10**). Thus, incorrect read-transcript assignment due to read truncation at least partially explains LR dropouts, which are attenuated by MPAQT when SR data are combined with LR data.

### MPAQT incorporates machine learning models of mRNA abundance to improve isoform-level quantification

Even after combining SR+LR data, transcript quantification remains challenging in certain contexts, exemplified by our single-cell benchmarking results (**Fig 4i-j**). Thus, we need additional sources of information, beyond sequencing reads, that further constrain the latent transcript abundances. Isoform expression is regulated by instructions encoded in the genome; thus, by estimating the regulatory logic underlying the transcript abundance profile of each given sample, we can further constrain the space of possible isoform abundances and potentially improve quantification accuracy. Sequence-to-expression models can potentially achieve this by learning, from (noisy) data, a sample-specific map connecting the genomic sequence features to mRNA abundance, which can then be integrated with empirical evidence from RNA-seq data to improve quantification.

This possibility is supported by our observation that isoform abundances can be predicted from the binding sites of sequence-specific RNA-binding proteins (RBPs). Specifically, we calculated the affinities^18^ of 128 RBPs, based on their known motifs^19^, toward the 5’ and 3’ untranslated regions (UTRs) of each mRNA isoform, removed redundancies by grouping motifs with similar affinity profiles into 35 motif archetypes, and, for each time point and replicate in our neuronal differentiation dataset, developed a random forest machine learning (ML) model that predicts each mRNA’s abundance from its motif archetype scores (see **Methods** for details). Based on gene-stratified five-fold cross-validation experiments, our ML models achieved an overall Pearson correlation of 0.38 (**Figure 6a**, minimum and maximum *r* of 0.36 and 0.40 across individual timepoints/replicates). Importantly, these models were able to capture patterns of differential isoform abundance across time points: for mRNAs with significant differential abundance across time points (one-way ANOVA), the sequence-based predictions correlated strongly with the observed differential abundances (mean r of 0.35 and 0.64 for isoforms with significant differential expression at FDR ≤0.05 and ≤0.01, respectively; **Figure 6b**). Analysis of the Shapley additive explanations (SHAP) suggests that our random forest ML models learn time point-specific rules about how RBP binding sites, especially at the 3’ UTR, are associated with isoform abundance (explored in **Supplementary Figure 11a**). Interestingly, we found that our models explain isoform usage patterns of many genes at each time point (**Supplementary Figure 11b** and **Figure 6c**)—specifically, for genes with more than two isoforms, differences in UTR sequence motifs predict isoform usage in differentiated neurons with mean *r*=0.23. For 25% of such genes (2508 out of 10,010), the correlation between UTR-based predictions and isoform usage in differentiated neurons exceeds 0.7; this fraction further increases to 41% if we focus on a subset of genes with highly reproducible isoform usage profiles (166 out of 405 genes with isoform imbalance F-value >500; **Figure 6c**). For genes with two isoforms and highly reproducible isoform imbalances (F-value >500), in 73% of cases (377 out of 517) the UTR-based predictions can correctly identify the dominant isoform.

**Figure 6.**
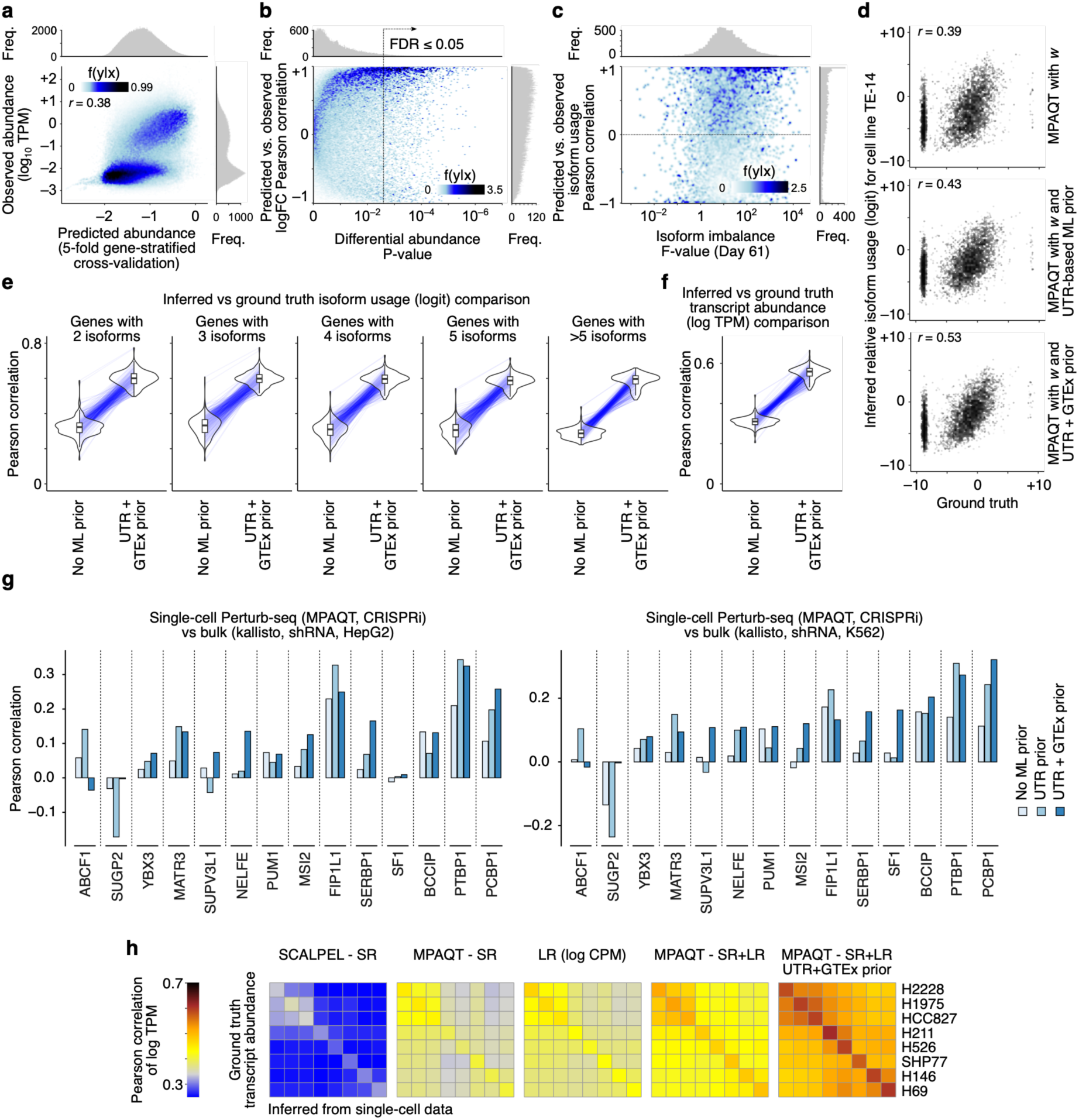
Integration of RNA-seq data with sequence-to-expression models for improved isoform quantification. (**a**) Observed vs. predicted transcript abundances in the neuronal differentiation dataset (5-fold gene-stratified cross-validation). Each point represents one transcript in one time point/replicate. Histograms represent marginal distribution of predicted (x-axis) and observed (y-axis) log-TPM values. (**b**) Correlation of predicted vs. observed log-fold changes during differentiation. Each point represents one transcript. The x-axis represents the statistical significance for differential log-TPM across time points (one-way ANOVA test). The y-axis represents the Pearson correlation between predicted and observed abundances across time points/replicates for each transcript. (**c**) Pearson correlation of predicted vs. observed relative isoform usage for each gene. Each point represents one gene with at least three isoforms. The x-axis shows the F-value from one-way ANOVA test for unequal abundances of isoforms. (**d**) Relative isoform usages inferred from pseudobulked single-cell data for TE-14 cell line (y-axis) vs ground truth values (x-axis, based on bulk RNA-seq) without integration with sequence-to-expression predictions (top), integration with UTR-based sequence-to-expression predictions (middle), or integration with UTR+GTEx-based sequence-to-expression predictions (bottom). Plotting details are similar to Figure 3c. (**e**) Systematic comparison of inferred vs ground truth relative isoform usages across 185 cancer cell lines in the CCLE mixture, similar to Figure 3d, but for pseudobulk single-cell MPAQT inferences without a sequence-to-expression prior and with UTR+GTEx-based sequence-to-expression priors. (**f**) Correlation between inferred vs ground truth transcript abundances in the CCLE mixture. (**g**) Comparison of log-fold changes inferred by MPAQT from CRISPRi-based single-cell Perturb-seq of RBPs vs log-fold changes inferred by kallisto from bulk data for shRNA-based knockdown of each RBP. Only RBPs with reproducible log-fold changes based on bulk data are included (Pearson correlation >0.05 between K562 and HepG2). (**h**) All-vs-all comparison of ground truth transcript abundances (rows) vs inferred transcript abundances (columns) in the LongBench dataset. Each inference approach is shown in a separate heatmap, with color gradient representing the correlation between inferred log-TPM and ground truth log-TPM across protein-coding transcripts. Data underlying this figure can be found in **Supplementary Data Table 6**.

Together, the above results suggest that even relatively simple sequence-to-expression models that only use UTR-RBP interactions can predict context- and isoform-specific expression. MPAQT can integrate these predictions with the empirical evidence from RNA-seq data to resolve ambiguities in assignment of reads to isoforms. Specifically, after initial estimation of isoform abundances from RNA-seq data by MPAQT, a sequence-to-expression model is built, whose predictions are then fed back to MPAQT in the form of a prior for subsequent rounds of inference from RNA-seq data (see **Methods** for details). **Fig 6d** shows an example, based on single-cell data from the CCLE cell line mixture, where incorporating UTR-based abundance predictions substantially improves isoform quantification from single-cell data. The correlation between the inferences and ground truth further improves if RBP-UTR interactions are augmented by isoform-specific low-dimensional embeddings learned from principal component analysis (PCA) of multi-tissue bulk RNA-seq data (**Fig 6d**, bottom; see **Methods** for details)—in this embedding space, isoforms that are co-expressed are embedded close to each other, allowing MPAQT to share information across isoforms that are likely under similar regulatory influences. These improvements are consistent across various cell lines, and can be seen both at the level of measuring relative isoform usage (**Fig 6e**) and absolute transcript abundance (**Fig 6f**).

We next asked whether these improvements translate into more accurate inference of transcript-specific regulatory responses. To this end, we performed CRISPR interference (CRISPRi)-based Perturb-seq targeting 33 RNA-binding proteins (RBPs) in K562 cells (see **Methods**), and compared transcript-level log-fold changes inferred from single-cell (pseudobulk) data with independent shRNA knockdown RNA-seq experiments from ENCODE performed in HepG2 and K562 cells^35^. These 32 RBPs were selected from previous work showing that they may regulate mRNA abundance indirectly through modulation of other mRNA stability factors^36^. We note that transcript-level perturbation responses vary substantially across experiments, cell lines, and/or perturbation modalities—for example, the correlation between RBP-matched perturbation profiles in the two ENCODE cell lines ranges from –0.11 to 0.26 (**Supplementary Figure 12a**). Thus, in our analyses, we focused on RBPs exhibiting reproducible transcript-level effects between the two ENCODE cell lines (ranging from Pearson correlation 0.05 for MSI2 to 0.26 for SUPV3L1), as well as transcripts whose responses were themselves reproducible across perturbations (see **Methods**). Within this subset, pseudobulk transcript-level log-fold changes inferred by MPAQT showed measurable agreement with the ENCODE perturbation profiles in both HepG2 and K562 cell lines despite differences in perturbation modality (**Fig 6g**). Importantly, incorporating sequence-to-expression priors consistently increased these correlations relative to inference without priors, with the largest improvements obtained by combining UTR-derived RBP binding features with PCA-based transcript embeddings (**Fig 6g**). These results suggest that sequence-informed priors not only improve agreement with baseline transcript abundance, but also enable more accurate inference of transcript-specific regulatory responses from scRNA-seq data.

To further validate this approach in an independent benchmark, we analyzed the LongBench dataset^34^, which provides matched bulk and single-cell transcriptomic measurements across multiple human cell lines as well as short- and long-read single-cell data. In this dataset, we obtained the most accurate isoform-level inferences from single-cell data when LR data were combined with SR data, SR positional biases were taken into account, and UTR features as well as PCA-based isoform embeddings were used to derive cell line- and isoform-specific priors (**Fig 6f**). In fact, with this combined approach, the correlation between isoform abundances inferred from single-cell data and the abundances inferred from bulk RNA-seq data (median *r*=0.6) reaches ∼78% of the correlation between simulated bulk RNA-seq technical replicates (median *r*=0.77; **Supplementary Figure 12b**). For comparison, SR-only inferences with SCALPEL achieve a median correlation of only 0.32, and LR-only data achieve a median correlation of 0.43, highlighting the substantial improvement that our integrative strategy provides relative to existing methods.

## Discussion

Transcript isoform quantification is fundamentally an inference problem, where the transcript abundances that we seek to estimate are latent variables that must be reconstructed from incomplete and noisy sequencing observations. Existing methods have largely approached this problem in the context of individual sequencing technologies, developing specialized algorithms for either short-^6^ or long-read ^37^ data. In contrast, MPAQT formulates transcript quantification as inference within a unified probabilistic generative framework, in which sequencing technologies are viewed as complementary measurement processes of the same underlying transcriptome. As we show in our results, this framework naturally enables joint analysis of heterogeneous modalities while remaining sufficiently flexible to incorporate platform-specific characteristics, sequencing biases, and biological priors in the form of sequence-to-expression models. These modeling capabilities are complemented by fast runtime and low memory requirements, especially in single-cell short-read analyses (**Supplementary Figure 13**). Specifically, in bulk data, MPAQT requires only modestly more runtime and peak memory relative to kallisto (**Supplementary Figure 13a-b**). In single-cell data, without ML prior, MPAQT is ∼13 times faster than SCALPEL with ∼4-fold less peak memory usage. The memory usage increases only slightly with the incorporation of ML prior, although the runtime increases to almost the same levels as SCALPEL, which can be rescued by refitting the ML prior periodically as opposed to in every iteration (**Supplementary Figure 13c-d**). Similar trends are observed for index construction (**Supplementary Figure 13e-f**). Such comparably light computational requirements make MPAQT particularly suitable for single-cell analyses in resource-limited settings.

MPAQT’s multi-platform integration can provide substantially more accurate transcript quantification than any single technology, as we demonstrate across a range of benchmarks in bulk and single-cell data. Short-read sequencing provides the depth required for precise estimation of transcript abundances across a wide dynamic range, but many reads cannot be uniquely assigned to individual isoforms. On the other hand, while largely resolving these ambiguities, long-read sequencing remains limited not only by lower sequencing depth, but also by experiment-specific technical biases and artifacts that cannot be overcome by simple depth scaling, ranging from nucleotide content biases to transcript length effects and artefacts created by read truncation. MPAQT exploits the fact that each technology constrains different aspects of the latent transcript abundance space to provide state-of-the-art transcript quantification. Importantly, for genes with complex splicing landscapes, we observed these improvements even when relatively modest amounts of long-read sequencing were incorporated (e.g., with as little as ∼200K simulated long reads, **Fig 4c**), suggesting that substantial gains in transcript quantification can already be achieved without requiring extremely deep long-read datasets. Importantly, we expect this general framework to remain applicable as RNA sequencing technologies continue to evolve, allowing future data types to be incorporated without fundamentally changing the statistical model.

This increased accuracy in isoform quantification also translates to downstream tasks, such as exon PSI inference from isoform abundances, as we show in our RT-PCR validation experiments (**Fig 4h**), further underlining the notion that isoform quantifications from SR+LR data are more accurate than isoform quantifications from SR data alone. We emphasize, however, that in this work, we used exon usage inference as an instrument to evaluate the accuracy of isoform-level inferences. Exon PSI can also be inferred from SR data based on direct measurement of reads supporting inclusion or exclusion of each exon—we observed slightly better agreement with RT-PCR results for our indirect exon usage inferences from SR+LR data than for direct PSI inferences obtained from rMATS^38^ (**Supplementary Figure 14**), but this comparison was limited to only four exons present in rMATS output. The extent to which this observation is generalizable, thus, requires further examination, and probably depends on a host of factors, including transcript annotation quality and completeness. In fact, our analyses suggest that transcript annotation can potentially affect isoform quantification: when we augment existing transcript annotations (from GENCODE^39^) with novel transcripts identified from our neuronal differentiation LR data (by Bambu^37^), we observe that the MPAQT-estimated abundances for 30-35% of previously annotated transcripts coming from genes with novel isoforms change by more than 10% (**Supplementary Figure 15**). Thus, incomplete annotation of transcripts confounds the quantification of other annotated isoforms from same genes.

Interestingly, MPAQT outperforms existing tools even for analysis of SR data alone, as we show in our SR-only benchmarking results (**Figure 2**). This improvement is specifically noticeable for a subset of transcripts that often appear as “dropouts” in other SR analysis tools. Anecdotally, we noticed that many of these dropouts and misquantified transcripts have “private” OUs that are not shared by other transcripts—when no reads are assigned to these private OUs, the overall transcript abundance is reported as zero by kallisto and Salmon, despite non-zero counts in other OUs (example shown in **Supplementary Figure 16a**). In other words, these tools over-emphasize the absence of reads in private OUs, even though a non-zero transcript abundance may lead to zero OU counts due to Poisson randomness. **Supplementary Figure 16b** demonstrates how this behaviour arises from the shape of the likelihood function: for kallisto, the maximum likelihood estimate (MLE) is at zero due to a monotonically decreasing likelihood function, even though non-zero abundances are also highly likely (note the relative plateau at the left end of the log-likelihood function). This leads to divergence of the MLE and the posterior mean. In contrast, for MPAQT, the mode of the likelihood function lies close to the posterior mean (**Supplementary Figure 16c**). Thus, MPAQT not only improves quantification by combining the complementary strengths of different platforms, but also by an improved statistical model for individual platforms.

The benefits of improved transcript quantification by combining SR and LR data are especially apparent when quantifying mRNAs from longer genes with complex alternative splicing landscapes, characteristics that are particularly enriched among genes involved in nervous system development. Of note, the improved transcript abundance estimates generated by SR+LR analysis in our *in vitro* model of neuronal differentiation enabled us to investigate sequence determinants of expression regulation at isoform resolution, suggesting that RBP recognition sequences located in 3’ UTRs are among the most important determinants of isoform abundance (**Supplementary Figure 11**). While not explored in our main results, we would like to note that, to our surprise, our UTR-based ML models can predict PSI of internal cassette exons with a Pearson correlation of 0.65 (**Supplementary Figure 17a-c**), and distinguish “included” exons (defined as those with PSI≥0.8) from “excluded” exons (PSI≤0.2) at AUROC (area under the receiver operating characteristic curve) of 0.92 (**Supplementary Figure 17d**), despite having no knowledge of the local sequence features of the cassette exon. These observations suggest that the sequence determinants of exon inclusion rate are not limited to the local context of the exon, and distal elements in the UTRs contribute significantly to the exon usage landscape, potentially through non-splicing mechanisms such as isoform-specific regulation of mRNA stability (**Supplementary Figure 11b**). A substantial body of work has been dedicated to identifying the determinants of cassette exon usage, including ML-based modelling of the sequences that surround these exons (e.g., see refs^27–30^). However, recent work has highlighted the challenges of predicting cell type-specific exon inclusion using only the local sequence features, especially in neurons^31^. Our results underline the importance of considering global mRNA sequence features in models of splicing regulation and exon usage, given that non-local sequence features can also affect exon inclusion levels through splicing-independent mechanisms, such as isoform-specific regulation of mRNA stability. Identification of such splicing-independent determinants of exon inclusion can also have implications in the design of therapeutics aimed at modulating disease-associated exons^32^.

Importantly, we show that, beyond revealing the sequence determinants of isoform regulation, such sequence-to-expression models can be readily incorporated into MPAQT as probabilistic priors that are continuously updated by empirical evidence from sequencing data to achieve substantial gains in quantification accuracy. Our results suggest that this approach particularly improves isoform quantification from single-cell data, where extreme 3’ (or 5’) biases severely limit the ability of SR data to resolve isoforms, which can only partially be alleviated by addition of LR data. Although we demonstrate this concept using relatively simple models based on RNA-binding protein motifs and low-dimensional transcript embeddings, the same framework is readily compatible with increasingly sophisticated sequence-to-expression models, including emerging RNA foundation models^23, 24^ capable of learning rich representations of transcript regulation. We anticipate that this interaction between mechanistic statistical inference and increasingly powerful sequence-based models will become an important direction for future transcriptomic analyses, allowing prior biological knowledge and experimental observations to reinforce one another within a unified probabilistic framework.

## Methods

### MPAQT generative model

As described in the **Results** section, MPAQT’s generative model connects the latent transcript abundances to the expected counts of a set of “observation units” (OUs), which are defined based on the technology/platform. Consider an RNA-seq dataset, generated by *K* different platforms from sequencing the same mixture of transcripts from the set *T*, with each transcript *t*∈*T* having the relative abundance *f_t_* so that Σ*_t∈T_ f_t_*=1. We also define ***f***=(*f*_1_*,…,f*_|*T*|_)^T^ to be the vector of relative abundances; ***f***∈(0,1)**^|^***^T^*^|^. For each platform *k*∈{1,…,*K*} and each transcript *t*, let’s define the “effective length^6^” *l_k,t_* to be a normalization factor such that *f_t_l_k,t_*=P*_k_*(*t*), where P*_k_*(*t*) is the probability of observing a read (or fragment) from transcript *t* in platform *k* (Σ*_t∈T_* P*_k_*(*t*)=1). In other words, P*_k_*(*t*) is the expected proportion of reads that originated from transcript *t*, given the transcript abundance profile and the platform.

Each read from each platform *k* is assigned to one OU from the set *U_k_*. For a read that originated from a given transcript *t*, the probability of being assigned to a given observation unit *u*∈*U_k_* is represented by P*_k_*(*u*|*t*). In other words, P*_k_*(*u*|*t*) is the probability of a read mapping to *u* conditional on that read having been selected from *t*. We also define *p_k,u,t_*=*l_t,k_*P*_k_*(*u*|*t*). Note that *p_k,u,t_* does not depend on the abundance of transcript *t*, and is rather a function of transcript properties/sequence and the platform *k*. It follows that P*_k_*(*u*) = Σ*_t∈T_* P*_k_*(*t*)P*_k_*(*u*|*t*) = Σ*_t∈T_ f_t_l_k,t_*P*_k_*(*u*|*t*) = Σ*_t∈T_ f_t_ p_k,u,t_*, where P*_k_*(*u*) is the probability that a read in platform *k* is assigned to the observation unit *u* (i.e., P*_k_*(*u*) is the expected proportion of reads in the dataset that map to *u*; Σ*_u_*_∈_*_Uk_* P*_k_*(*u*)=1). Subsequently, the expected number of reads from platform *k* mapping to *u* is given by *λ_k_*_,*u*_= P*_k_*(*u*)*N_k_*, where *N_k_* is the total number of reads obtained from platform *k*. In turn, the observed number of reads from platform *k* mapping to *u* is drawn from a Poisson distribution (which is commonly used to model count data, e.g., see ref^40^), with expectation *λ_k_*_,*u*_. This generative model can be summarized as follows:

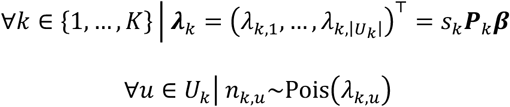

Here, ***P****_k_*∈**ℝ**_≥0_**^|^***^Uk^*^|×|*T*|^ is a platform-specific matrix whose elements, *p_k,u,t_*, are described above, ***β***∈**ℝ**_+_**^|^***^T^*^|^ is a column vector of scaled abundances for transcripts *T*, and *s_k_* is a platform-specific scaling factor. We also impose an isotropic Gaussian prior on log of ***β***:

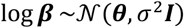

Here, ***θ***∈**ℝ^|^***^T^*^|^ represents transcript-specific means of the prior. In the absence of an informative prior, we use ***θ***=**0** to enforce a unique solution (since ***β*** and *s_k_* together form an underdetermined system, given that ***β*** and *s_k_* are differently scaled representations of ***f*** and *N_k_*, respectively, so that ***β***=*N**f/**s_k_*). As we discussed in the Discussion section, this prior also prevents dropouts by closing the gap between the MAP estimate and the posterior mean. In the presence of an informative prior (e.g., predictions by a sequence-to-expression model), ***θ*** is set to the vector of log-abundances predicted by the model.

Given {***n***_1_, ***n***_2_, …, ***n****_K_*}, which is the set of observed OU counts across *K* platforms, and {***P***_1_, ***P***_2_, …, ***P****_K_*}, which is the set of matrices that connect transcript identities to OU probabilities in each platform as described above, MPAQT finds the maximum a posteriori (MAP) estimate of ***β*** and {*s*_1_, *s*_2_, …, *s_K_*} using an expectation maximization (EM) algorithm, as described in **Supplementary Methods**. The MAP estimate of ***β*** is then used to obtain the vector of relative abundances ***f*** (and TPM) by rescaling.

### Obtaining the platform-specific matrices *P_k_*

For any platform *k* that represents short-read (SR) data, we obtain the matrix ***P****_k_* by enumerating all possible reads from a reference transcriptome in which all transcripts have exactly equal abundances. Specifically, given read length *l_r_*, MPAQT first appends poly(A) tails of length *l_r_*–*k*–1 to transcript biotypes that are commonly poly-adenylated (protein-coding, lncRNA, and other biotypes excluding small RNAs), followed by generating every possible read of length *l_r_* at stride 1. Here, *k* is the length of *k*-mers used by kallisto for indexing; thus, a poly(A) length of *l_r_*–*k*–1 ensures that reads overlapping the poly(A) tail have at least *k* informative nucleotides for mapping to the transcriptome. The generated reads are then mapped back to the transcriptome and assigned to ECs using kallisto^6^. Since each generated read is tagged with its transcript of origin (*t*) and is mapped to a unique EC (*u*), we can calculate the proportion of reads that originate from transcript *t* and map to EC *u*, i.e., *p_u,t_*. In practice, however, we do not calculate the proportions *p_u,t_*, but instead directly use the read count *m_u,t_*. Since *m_u,t_* is proportional to *p_u,t_*, it only affects the scale *s_k_*.

For long-read data, in the simplest scenario, we can assume that each long read is unambiguously assigned to one transcript; thus, *U_k_*=*T*, and P*_k_*(*u*|*t*)=1 when *u*=*t* and zero otherwise. Furthermore, we can assume that read counts are proportional to the transcript abundances, i.e., no biases exist and all transcripts have the same effective length (∀*t*∈*T l_k_*_,*t*_=1). The matrix ***P****_k_* for long-read sequencing data is then simply the |*T*|×|*T*| identity matrix ***I***. However, both these assumptions can be violated in real-life applications. While the unambiguity assumption may be true for a large fraction of long reads, factors such as transcript degradation, read truncation, and other sequencing or alignment artifacts can introduce ambiguities in at least a fraction of read-transcript assignments^37^. The statistical framework of MPAQT in principle allows for these ambiguities to be taken into consideration, for example by using “read classes”^37^ as OUs, each of which may be compatible with multiple isoforms. The current version of MPAQT, however, is based on assumption of unambiguity. Regarding the second assumption, as discussed in the Results section, we have found that substantial length and GC-biases exist in long-read data. Therefore, MPAQT provides the option to explicitly learn these biases from data and incorporate them in the matrix ***P****_k_*. We model *l_k_*_,*t*_, the effective length of transcript *t* in long-read platform *k*, as a function of transcript-level variables ***c****_t_* (***c****_t_* ∈**ℝ**^|*D*|^, where *D* is the set of transcript-level covariates whose effects we want to model). Specifically, log *l_k_*_,*t*_ = ***c****_t_***⋅*γ****_k_*, where **⋅** is the dot product, and ***γ****_k_*∈**ℝ**^|*D*|^ is the vector of coefficients representing the effect of the covariates on the propensity of the transcripts to be captured by long-read platform *k*. The log-link ensures that *l_k_*_,*t*_ is restricted to the domain **ℝ** ^|*T*|^. The MAP estimate of ***γ****_k_* is obtained during model fitting as described in **Supplementary Methods**.

### Positional bias correction

Single-cell RNA-seq protocols exhibit extreme positional biases (e.g., 3’ enrichment in Chromium Single Cell 3’ kits). MPAQT corrects this by learning positional weights from pseudobulk of all cells before quantifying abundances at the level of cell types/clusters. During generation of matrix ***P****_k_* (indexing), MPAQT records, for each read that maps to a given OU *u*, its distance from the 3’ (or 5’) end of the transcript *t* that generated the read. It then averages these distances for every OU-transcript pair (*u*,*t*) to obtain *d_u_*_,*t*_, the average distance from transcript end for reads originating from transcript *t* and mapped to OU *u*. Consider the positional weight function *w_k_*(*d*), representing the capture efficiency in platform *k* for reads at distance *d* from transcript end. For reads that originate from transcript *t* and map to OU *u*, the mean capture efficiency can be approximated as *w_k_*(*d_u_*_,*t*_). Thus, MPAQT modifies element *p_k,u,t_* of matrix ***P****_k_* by multiplying it by *w_k_*(*d_u_*_,*t*_). In other words, ***P****’_k_ =*[*p’_k,u,t_*] replaces ***P****_k_* in the generative model, where *p’_k,u,t_*=*w_k_*(*d_u_*_,*t*_)*p_k,u,t_*:

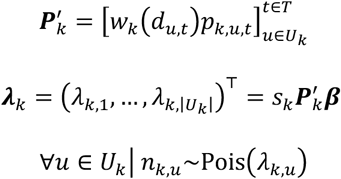

MPAQT models *w_k_*(*d*) as a piecewise constant function over *D* intervals:

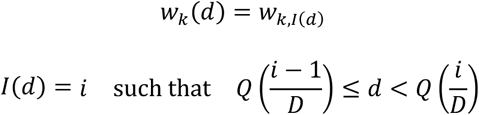

Here, *Q* is the quantile function. Thus, *i*∈{1,…,*D*) indexes intervals that represent *D* equally spaced quantiles over all *d_u_*_,*t*_ values. Starting with *w_k,i_*=1 for all *i*∈{1,…,*D*), these weights are optimized via an iterative expectation-maximization procedure where, in each iteration, given the current *w_k_*_,*i*_ values, transcript abundances ***β*** and library size *s_k_* are estimated, followed by L-BFGS-B optimization of all *w_k_*_,*i*_ values given the MAP estimate of ***β*** to maximize the likelihood of OU counts. Once *w_k_*_,*i*_ values are optimized, they are kept frozen and the resulting ***P****’_k_* is used to estimate ***β*** in each cell type/cluster separately.

### Processing of the Quartet dataset

The Quartet dataset was downloaded from CNCB-NGDC (https://download.cncb.ac.cn/gsa-human/HRA001859/) and were mapped to the data and metadata on the Chinese Quartet portal using md5sums (https://chinese-quartet.org/#/data/download/quartet-transcriptomics). Two high-quality batches, Batch3 (P_ILM_L1_B1) and Batch21 (R_ILM_L8_B1), were used for this analysis (Supplementary Table 3 of ref^25^). Batch 3 (P_ILM_L1_B1) consists of the following CNCB-NGDC sample identifiers: D5 (HRR777581, HRR777582, HRR777583), D6 (HRR777584, HRR777585, HRR777586), F7 (HRR777587, HRR777588, HRR777589), and M8 (HRR777590, HRR777591, HRR777592). Batch 21 (R_ILM_L8_B1) consists of the following CNCB-NGDC sample identifiers: D5 (HRR777669, HRR777670, HRR777671), D6 (HRR777672, HRR777673, HRR777674), F7 (HRR777675, HRR777676, HRR777677), and M8 (HRR777678, HRR777679, HRR777680). Sequencing run metadata was mapped from the sequencing dates (P_ILM_L1_B1: 20200602, R_ILM_L8_B1: 20180928) (based on Supplementary Table 2 of ref^25^). Sequencing dates, which were unique among all 21 batches, were then used to identify the corresponding batches on the Chinese Quartet data portal, where the sequencing dates are present in the file names of the data. The qRT-PCR and ddPCR fold-change data were downloaded from the Quartet paper (Supplementary Table 9 of ref^25^). Kallisto (v0.50.1), salmon (v1.3.0), RSEM (v1.3.3) and MPAQT-SR were run on all 24 samples, and gene-level quantifications from six total replicates (two batches of three) were summarized in two steps. First, for each gene and tool, the mean TPM was calculated across the three replicates within each batch. The resulting batch means were then averaged to obtain a single overall mean TPM per gene across both batches. Log fold-change (logFC) values were subsequently calculated to compare expression between conditions by computing log_2_ ratios for the following contrasts: D5 vs D6, F7 vs D6, and M8 vs D6. Gene-level benchmarking was then performed for each tool, combining all four contrasts into a single plot, totalling 91 gene-level comparisons per tool (**Figure 1b**).

### Processing of the MAQC dataset

The MAQC data was taken from GEO accession GSE83402, and single-end samples MAQCA_1 (four technical replicates: SRR3670977, SRR3670978, SRR3670979, SRR3670980) and MAQCB_1 (four technical replicates: SRR3670985, SRR3670986, SRR3670987, SRR3670988) were processed with RSEM^13^ (version 1.3.3, following alignment with bowtie2 version 2.4.2), salmon^12^ (version 1.3.0, with the --validateMappings and --gcBias flags), kallisto^6^ (version 0.48.0) and MPAQT.. Technical replicates for each sample were combined (at the level of FASTQ files) during quantification. Differential expression was calculated as the logarithm of fold-change (logFC) between MAQCB_1 and MAQCA_1, separately for each quantification tool.

### Generation and processing of simulated datasets

Simulated datasets for benchmarking, in the form of paired-end FASTQ files, were generated from ground truth TPM values using the simReads function from the Rsubread R package^35^. Rsubread takes as input the number of reads to simulate and a list of transcripts with their desired TPMs. We generated two simulated datasets using two different sets of ground truth TPMs. For the first dataset, ground truth TPMs were sampled from an exponential distribution (using rexp R function with default rate=1). For the second dataset, we first used kallisto to quantify transcript abundances from RNA-seq data of the MDA-MB-231 cancer cell line (GEO entries GSM4886854, GSM4886855)^37^ and then used the resulting TPMs as the ground truth for read simulation. For the rexp.sim dataset, three simulated “replicates” were generated, and one sample was generated for the MDA-MB-231-based dataset, each with 30 million paired-end reads of 75 bp. These samples were processed with RSEM^13^ (version 1.3.3, following alignment with bowtie2 version 2.4.2), salmon^12^ (version 1.3.0, with the --validateMappings and --gcBias flags), kallisto^6^ (version 0.48.0) and MPAQT..

### Processing of the Sequins spike-in dataset

Illumina short-read sequencing data were downloaded from the Sequence Read Archive with the following identifiers: SRR14286057 (NCI-H1975_R1), SRR14286058 (NCI-H1975_R2), SRR14286059 (NCI-H1975_R3), SRR14286066 (HCC827_R1), SRR31761441 (HCC827_R2_1), SRR31761442 (HCC827_R2_2), and SRR14286069 (HCC827_R3). For sample HCC827_R2, two technical replicates were provided in the data (SRR31761441 and SRR31761442), which were combined into a single sample. ONT long-read sequencing data were downloaded from the Sequence Read Archive with the following identifiers: SRR14286054 (R1_H1975_ONT), SRR14286055 (R2_H1975_ONT), SRR14286056 (R3_H1975_ONT), SRR14286063 (R1_HCC827_ONT), SRR14286064 (R2_HCC827_ONT), and SRR14286065 (R3_HCC827_ONT). PacBio long-read sequencing data were downloaded from the Sequence Read Archive with the following identifiers: HCC827_R1 (SRR14286070), HCC827_R2 (SRR14286071, SRR14286072), HCC827_R3 (SRR14286073, SRR14286074), NCI-H1975_R1 (SRR14286060), NCI-H1975_R2 (SRR14286061), and NCI-H1975_R3 (SRR14286062). Samples with more than one corresponding data file were combined into a single FASTQ file before processing. Sequins data and metadata, including expression tables for Mix A and Mix B, GTF annotation file, and FASTQ sequences, were downloaded from https://github.com/XueyiDong/LongReadRNA.

Long-read RNA-seq data was processed as follows: minimap2 (v2.26) was used to map reads to reference genome GRCh38, then Bambu (v3.8.3) used to quantify samples. Bambu output was then benchmarked against ground truth or used as input to MPAQT-SR+LR.

### Processing of UHRR samples

Illumina HiSeq 2000 short-read samples from the Sequencing Quality Control (SEQC) project were downloaded from SRA project PRJNA208369 for UHRR (Sample A)^33^, consisting of two samples from each of two sequencing centers (NYG: SRR1215997, SRR1216019; BGI: SRR896666, SRR896676). ONT data for UHRR was downloaded from SRR12010483^41^, and PacBio data from the Pacific Biosciences Sequel II System Data Release [https://github.com/PacificBiosciences/DevNet/wiki/Sequel-II-System-Data-Release:-Universal-Human-Reference-(UHR)-Iso-Seq].

TaqMan RT-PCR measurements were downloaded from GSE5350 (GSM129638, GSM129639, GSM129640, GSM129641), and mean expression across the 4 replicates were calculated. The metadata was obtained from GPL4097. RefSeq transcript IDs (prefixed with “NM”, indicating they are protein coding mRNAs) were mapped to GENCODE transcript IDs using biomaRt::getBM. RefSeq IDs that did not map to a GENCODE transcript, or that mapped to more than one GENCODE transcript, were filtered out, leaving 906 TaqMan RT-PCR measurements.

For short-read analysis of UHRR samples, kallisto (v0.50.1), salmon (v1.3.0), and MPAQT-SR were run on all four samples using GENCODE v47 annotations, and then the mean TPM predictions were calculated across the four samples. For MPAQT joint short-read and long-read analysis of UHRR samples, long-read data was aligned to reference genome GRCh38 using minimap2 (v2.28), with flags *-x map-ont* (for ONT) and *-x map-hifi* (for PacBio), followed by quantification with Bambu (v3.8.3) with GENCODE v47 GTF. MPAQT was then run using both Bambu output and Illumina short-read data as input. MiniQuant (v1.4) using joint SR and LR mode was run on each of the four Illumina samples, along with either ONT or PacBio data, specified using either the the *dRNA-ONT* or *cDNA-PacBio* options for the *--long_reads_library_prep* flag, respectively.

### Cortical neuron differentiation and RNA-seq data generation

#### Cortical neuron differentiation

The hESC SOX10::GFP bacterial artificial chromosome reporter line (in the H9 background) was used for neural differentiation according to the protocol adapted from a study of brain organoids^36^. In brief, the hESC line was maintained in feeder-free conditions with the E8 medium. Neural differentiation was initiated when the cells reached 90-100% confluency. From days 0-11, the cells were maintained in neural induction medium (10 µM SB431542 and 100 nM LDN193189 in E6 medium) with medium change every two days. From day 12, the cells were fed the cortical neuron medium (10 ng/mL GDNF, 100 µM ascorbic acid, 1x Glutagro, 1x N2 supplement, 1x B27 without vitamin A in neurobasal medium) with medium change every other day until rosette structures became visible. Then, neurons were detached using Accutase and replated on poly-L-ornithin/fibronection/laminin-coated plates. Neurons were maintained in cortical neuron medium with medium change every other day. On days 22-24, neurons were checked for the presence of axonal projections and 10 µM DAPT was included in the cortical neuron medium until the projections appeared. From day 30, neurons were considered mature with the medium feeding frequency reduced to 1-2 times per week.

#### RNA extraction, short-/long-read RNA-seq library prep, and sequencing

Cells were harvested at days 0, 41, and 61, followed by RNA extraction using Zymo Quick-RNA Microprep kit according to the manufacturer’s protocol. SIRV set 4 (Lexogen) was spiked at 1% in the hESC and differentiated neuron-derived RNA samples. Short-read RNA-seq libraries were prepared using the SMARTer Stranded Total RNA-Seq Kit v3. Libraries were sequenced on a NextSeq 550 sequencer (2×75 bp paired-end). PacBio Iso-seq libraries from the same RNA samples were generated using the NEBNext Single Cell/Low Input cDNA Synthesis & Amplification Module, PacBio Iso-Seq Express Oligo Kit and SMRTbell express template prep kit 2.0 according to the manufacturer’s protocol. The libraries were sequenced on a PacBio Sequel IIe.

### Processing of short-read RNA-seq neuronal differentiation data

SR data were processed using RSEM^13^ (version 1.3.3, following alignment with bowtie2 version 2.4.2), salmon^12^ (version 1.3.0, with the --validateMappings and --gcBias flags), kallisto^6^ (version 0.48.0) and MPAQT.

SR sequencing data for the neuronal differentiation samples were processed using the paired-end options for the above tools. For this dataset, we added the spike-ins to the reference transcriptome of each tool to enable their quantification. Each spike-in was added in as its own separate chromosome, and 1000 “N” spacer nucleotides were added on either side of each spike-in sequence. As described above, we used the SIRV-Set 4 from Lexogen, which contains 114 spike-in transcripts. We used 107 in this analysis, since SIRV-403 to SIRV-410 were not included in the reference FASTA provided by Lexogen.

### Processing of long-read RNA-seq neuronal differentiation data

The IsoSeq pipeline (Pacific Biosciences) was used to process the neuronal differentiation LR data and generate circular consensus sequence (CCS) reads, which were stored in uBAM (unaligned BAM) format. Next, lima (PacBio) was used to remove primer sequences. IsoSeq3 ‘refine’ command was used to remove poly-A tails and concatemers (reads which are attached end-to-end), followed by the ‘cluster’ command to cluster reads that represent the same transcript (i.e. make them adjacent). The ‘align’ command of pbmm2 (PacBio) was then used to align reads to the reference genome, followed by the Isoseq3 ‘collapse’ command to condense the data into a transcriptome (fasta + GFF) and provide an abundance file containing full length counts (FL counts).

The quality control script from SQANTI3^38^ (sqanti3_qc.py) was used together with supporting data types, including CAGE peak, polyA motif list, polyA peaks file, and Intropolis splice junctions (https://github.com/nellore/intropolis), to remove low-quality transcripts according to SQANTI’s quality criteria. Next, the rules filter script (sqanti3_RulesFilter.py) was run to further filter transcripts based on the following criteria: if a transcript is a full splice match (FSM), then it is kept unless the 3’ end is unreliable (intrapriming); if a transcript is not a FSM, then it is kept only if all of below are true: (a) 3’ end is reliable; (b) the transcript does not have a junction that is labeled as RT Switching; and (c) all junctions are canonical. Finally, the full-length (FL) LR counts from SQANTI3 output that had the same “associated_transcript” were combined, providing transcript counts for input to MPAQT and for use in benchmarking.

For all analyses, reference transcriptome and genome annotations from GENCODE^39^ (v38 or above as stated in the sections below) was used, corresponding to human genome assembly GRCh38.p13.

### RT-qPCR measurement of differential cassette exon inclusion

#### Selection of cassette exons

To aggregate isoform-level abundances into cassette exon-level measurements, we obtained the annotation of cassette exons using the “generateEvents” command from SUPPA2 v.2.3^40^ for the reference transcript annotations. The abundances of all transcripts supporting each of the two possible outcomes of every event were then aggregated, providing, for each cassette exon in each sample, the sum-TPM of isoforms supporting the inclusion of the cassette exon and the sum TPM of isoforms supporting exon exclusion. This process was repeated separately for TPM inferences obtained by MPAQT using SR data and MPAQT using SR+LR data. We then fitted a limma^41^ model (using “limma” package v3.56.2) with the following formula for each cassette exon across all samples/measurement types: *y*∼*s*+*w*+*x*+*t*:*x*+*x:w*+*t*:*x*:*w*. Here, *y* is the log-sum-TPM, *s* is a multi-level sample indicator, *x* is a binary indicator of whether the measurement belongs to the inclusion (*x*=1) or exclusion (*x*=0) set of transcripts, *w* is a binary indicator of whether the measurement is based on SR+LR data (*x*=1) or SR-only data (*x*=0), and *t* is a multi-level timepoint indicator, with *t*=0 as the reference level. The coefficient of the *t*:*x*:*w* interaction term indicates the degree to which Δlogit-PSI between days 0 and 61 changes if we switch from SR-only measurements to SR+LR measurements (note that logit-PSI of each exon is equal to log-sum-TPM of inclusion minus log-sum-TPM of exclusion transcripts). We used the empirical Bayes functionality of limma to extract the coefficient and associated P-value of this coefficient, followed by selection of the top nine cassette exons with the smallest P-values (P<4×10^-5^).

#### RT-qPCR measurements

Transcript levels were measured using RT–qPCR by reverse transcribing total RNA to complementary DNA (Maxima H Minus RT, Thermo), then using PerfeCTa SYBR Green SuperMix (QuantaBio) per the manufacturer’s instructions (for primer sequences, see **Supplementary Data Table 4**).

### Processing of CCLE dataset

The short read (Illumina NextSeq 500) single-cell RNA-seq (10x Chromium 3 kit v2 library) data from CCLE was downloaded from GEO accession GSE157220^27^. The metadata file was taken from Broad Institute’s Single-cell portal (https://singlecell.broadinstitute.org/single_cell/study/SCP542/pan-cancer-cell-line-heterogeneity=study-download). The pool composition file was manually annotated with SRR accession numbers of corresponding bulk RNA-seq (Illumina HiSeq 2500) data, which were taken from PRJNA523380^42^. Bulk transcript quantification of all cell lines was performed using kallisto (v0.50.1) and MPAQT. For single-cell RNA-seq with MPAQT, positional bias was first inferred for each sequence-run. Single-cell transcript quantification of all cell lines was performed using MPAQT and SCALPEL at pseudobulk level using barcode annotations provided in the metadata. Estimated transcript counts by SCALPEL were converted to TPM. Missing transcripts in SCALPEL’s output were filled with 0. Gene quantification was obtained by aggregating abundances of all transcripts of the gene for kallisto, MPAQT, and SCALPEL outputs, and additionally obtained using STARsolo (v2.7.11a) at pseudobulk level for single-cell RNA-seq data. Gene counts generated by STARsolo were again converted to TPM. To benchmark MPAQT’s quantification output, we computed Pearson correlations between the estimated abundance using 4 versions of MPAQT (without positional bias, with positional bias, positional bias + UTR prior, positional bias + UTR_GTEX prior, as discussed below) and estimated abundances using kallisto, STARsolo and SCALPEL at 3 levels: transcript log-TPM, gene log-TPM and relative transcript usage—relative usage for each transcript was computed as its TPM divided by the TPM of its gene, followed by logit transformation.

### CRISPRi Perturb-seq of RNA-binding Proteins and data processing

The 32 RNA-binding proteins (RBPs) tested in this screen encompass: AATF, ABCF1, AKAP1, BCCIP, DDX55, DKC1, DNAJC2, EIF4B, FAM120A, FIP1L1, IGF2BP1, IGF2BP2, ILF2, MATR3, MSI2, NELFE, PABPC4, PCBP1, PTBP1, PUM1, QKI, RECQL, RPLP0, RRP9, RTF1, SERBP1, SF1, STIP1, SUCLG1, SUGP2, SUPV3L1, YBX3. With use of the dual-guide system, two sgRNAs per RBP, selected from the hCRISPRi-v3^43^, were individually cloned into the pJR85 plasmid (Addgene, 140095) via Golden-Gate ligation with BstXI, BlpI and BsmBI-v2, receiving a fragment from pJR89 (Addgene, 140096). The resultant plasmid library with the addition of 10 negative control plasmids generated in parallel was pooled in an equimolar ratio. Lentivirus was prepared in HEK293T cells (ATCC CRL-3216) using the TransIT-Lenti Transfection Reagent. K562 cells stably expressing Zim3-dCas9 (Addgene, 188766) were transduced with the library at low multiplicity of infection, yielding 5% transduction rate, sorted for BFP positivity after 72 h, and 300,000 cells were collected for sequencing. Four lanes of the Chromium Next GEM Single Cell 3’ HT Reagent Kit v3.1 (Dual Index) with Feature Barcode technology for CRISPR Screening (10x Genomics) were run and gene expression as well as sgRNA capture library preparation was performed according to the manufacturer’s instructions. Sequencing was performed on a NovaSeq X (Illumina).

Cell Ranger 7.0.0 (10x Genomics) was used to align scRNA-seq reads to the transcriptome and align sgRNA reads to the library, collapsing reads to UMI counts, and cell calling. Geomux^44^ was used to assign sgRNAs to cells, followed by forming pseudobulk for each RBP (and the non-targeting control group) based on cells assigned to the corresponding sgRNAs—the size of the pseudobulk groups ranged from 1250 to 4153 cells. Each of the four captures was processed separately, providing four replicates per perturbation. Single-cell transcript quantification was performed at the pseudobulk level for each RBP treatment and non-targeting control using MPAQT (with GENCODE v44 transcriptome) with positional bias modeling, either without an informative prior, with UTR-based prior, or with UTR+GTEx prior. For each variation of MPAQT, pseudobulk log-TPMs were averaged by treatment across the four libraries, followed by isoform-wise centering to obtain log-fold changes relative to the mean of all pseudobulks.

For bulk reference measurements, the raw FASTQ RNA-seq files of two cell lines (K562, HepG2) receiving 66 independent shRNA knockdown treatments targeting 32 RBPs and 27 control treatments were downloaded from ENCODE (https://www.encodeproject.org/files/). The data comprised 186 biological replicate samples. Isoform-level abundance for each sample was estimated using kallisto (v0.50.1) quant with the GENCODE v44 transcriptome. Only transcripts with TPM ≥ 1 in at least 5% of bulk samples were retained (95514 out of 252855). Log-TPM values were adjusted for potential batch effects using surrogate variable analysis (SVA^45^); the number of surrogate variables was estimated to be 16 using the Buja–Eyuboglu permutation method (20 permutations). Adjusted abundances were averaged across two biological replicates for each treatment, followed by isoform-wise centering to obtain log-fold changes relative to the mean of all samples (separately for K562 and HepG2). Bulk across-cell-line reproducibility was calculated using Pearson correlation between the K562 and HepG2 profiles. 14 out of 32 RBP treatments with Pearson correlation ≥0.05 (across transcripts) and 447 out of 95514 transcripts with Pearson correlation ≥0.6 (across RBP treatments) were retained. Pairwise Pearson correlations between Perturb-seq and bulk RBP knockdown profiles were calculated separately for K562 and HepG2.

### Processing of LongBench single cell data

The LongBench multi-platform reference dataset^34^, which contains bulk and single-cell data from eight cancer cell lines, were downloaded from GEO with (GSE303762). Bulk RNA-seq data for the eight cell lines were obtained from SRR30947507 (H1975), SRR30947506 (HCC827), SRR30947495 (H2228), SRR30947483 (H146), SRR30947474 (H69), SRR30947472 (SHP77), SRR30947473 (H526), and SRR30947477 (H211). Single-cell data for the cell line mixture from eight cancer cell lines were downloaded from SRR30947481 (Illumina short-read), SRR30947476 (PacBio long-read), and SRR30947479 (ONT long-read). GENCODE v44 was used for all analyses. ONT long-read counts provided by the authors were obtained from GSM9135504 (file GSM9135504_ont_sc_transcript_count_matrix.mtx.gz) and was used as input to MPAQT.

Cell barcode cell type annotations were kindly provided by the authors (**Supplementary Data Table 5**). The barcode-cluster annotation file contains 3,850 cells distributed across eight clusters: H146 (424), H1975 (704), H211 (318), H2228 (792), H526 (234), H69 (414), HCC827 (500), and SHP77 (464). For SCALPEL benchmarking, the LongBench short-read single-cell data was first split into 8 cluster-specific FASTQ files using the author-provided barcode-cluster annotation file. Of 117,515,267 total reads, 84,993,328 were assigned to clusters: H146 (4,900,034), H1975 (21,076,438), H211 (4,108,512), H2228 (22,134,232), H526 (3,481,095), H69 (5,029,416), HCC827 (16,615,059), and SHP77 (7,648,542), while 32,521,939 reads with barcodes absent in the metadata remained unassigned. Cellranger v7.1.0 was run on each cluster separately, and then SCALPEL^15^ (github commit 3c31ffc) was run on each cluster, using the apptainer image provided by the SCALPEL authors on Zenodo (SCALPEL.container.sif). Counts for each cluster were summed to produce cluster-level counts for each transcript, and raw counts were converted to TPM.

### Sequence-based prediction of mRNA abundances

#### RNA-binding protein (RBP) motifs

Human RBP motifs were downloaded from CISBP-RNA^19^ (v0.6) and filtered to include only motifs obtained by RNAcompete assays, encompassing 99 direct and 116 indirect (homology-based) motif-RBP associations, with 128 unique motifs and 128 unique RBPs (**Supplementary Data Table 6**). We used AffiMx^18^ to scan the 5’ and 3’ UTRs of all isoforms in GENCODE v38 with the 128 RNAcompete motifs, limiting to transcript isoforms with both annotated 5’ and 3’ UTRs. For each of the 5’ and 3’ UTR sets, the affinities were log-transformed and scaled for each motif separately (mean=0, variance=1), followed by application of convex nonnegative matrix factorization^42^ to obtain a non-redundant set of 35 “motif archetypes” and the score of each UTR for each archetype—each motif archetype is a convex combination of several motifs (usually with similar affinity profiles across the UTRs); in turn, the affinity profile of each motif can be reconstructed by a convex combination of motif archetypes. The number of motif archetypes was selected so as to result in a minimum Pearson correlation of 0.9 between the original and reconstructed affinity profiles per motif.

#### Training/validation of sequence-to-expression models and incorporation as prior in MPAQT

The 5’ and 3’ UTR motif archetypes were collated to obtain 70 motif archetype scores per transcript, which were used as predictive features for construction of machine learning models of isoform abundances. For modeling neuronal differentiation data, for each neuronal differentiation time point (days 0, 41, and 61) and each of the two replicates, random forest models were constructed to predict log10 TPM of each isoform from its 5’ and 3’ UTR motif archetype scores, using a 5-fold gene-stratified cross-validation approach. In other words, genes were randomly assigned to five different folds, each time all isoforms of the genes in one of the folds were held out, a random forest model was trained on the remaining isoforms (using R “ranger” package v0.16.0 with default parameters), and the model was used to predict the abundances of held-out transcripts.

To use sequence-to-expression model predictions as priors in MPAQT, we also augmented RBP motif archetype scores with principal component analysis (PCA) embeddings from analysis of multi-tissue bulk RNA-seq data from the Genotype-Tissue Expression (GTEx) Project^46^. Bulk RNA-seq transcript-level expression data were downloaded from GTEx (v8 release; file: GTEx_Analysis_2017-06-05_v8_RSEMv1.3.0_transcript_tpm.gct.gz; https://www.gtexportal.org/home/downloads/adult-gtex/bulk_tissue_expression), corresponding to GENCODE v44. A transcript-by-sample expression matrix was constructed by TPM values for all samples (dimension 244940×19789). Expression values were log-transformed (after adding a pseudocount of 10^−30^), standardized per sample (mean=0, variance=1), and PCA was performed (prcomp in R), treating transcripts as observations and samples as variables. The top K=200 principal components were retained, and the corresponding score matrix was extracted, where each row represents a transcript and each column corresponds to a principal component. These scores were subsequently concatenated with standardized RBP motif archetype scores to form feature matrix ***X*** (columns were normalized to have mean zero, followed by replacing missing values with 0), and used in a mixed-effects model of the form:

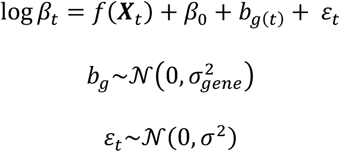

Here, *β*_0_ is a fixed-effect intercept, *b_g_* is a gene-specific random effect, and *ε_t_* is the residual term. The random effect intercept allows the model to consider gene-level determinants not included in the feature matrix (such as transcriptional effects). This model is iteratively updated at inference time by MPAQT, where, in each iteration, current transcript abundance estimates are used to fit the mixed-effect model, and the resulting transcript-level mean predictions (along with the fitted *σ*^2^) are used in the next iteration as prior. For the fixed-effect component *f*(***X****_t_*), we experimented with both random forests and linear models, and observed similar predictive performances, while linear models were substantially faster. Thus, MPAQT by default uses a linear mixed-effect model (gpboost in R).

### Benchmarking computational requirements

Three bulk RNA-seq libraries SRR8615406, SRR8616102, and SRR8615880 from CCLE were arbitrarily chosen and quantified using MPAQT and kallisto (v0.52.0). GENCODE v47 was used as the reference. Single-cell RNA-seq data from SRR12557905 were grouped into 24 cell-line clusters using provided barcode annotations and quantified against GENCODE v44 using MPAQT and SCALPEL. MPAQT was run with the default log-normal prior (no additional sequence informed prior) and with GTEx prior, with the latter refitted every iteration (k=1) or every 20 iterations (k=20). Between refits, the previous fitted prior was retained. SCALPEL was run separately for each cluster, with its internal salmon transcriptome indexing included in each run.

All quantification runs were performed on Intel Xeon Platinum 8468 nodes using 8 threads per tool. Each analysis was repeated three times, and the mean and standard deviation were calculated for wall-clock runtime and whole-process-tree maximum resident set size. Bulk runtimes were measured end-to-end for each sample, including pseudoalignment. Single-cell runtimes were measured end-to-end for the dataset, with cumulative wall time summed across the 24 clusters. Index construction was evaluated separately and excluded from these runtimes. Indexes were built using 16 threads on the same node type, with three runs per tool, using GENCODE v47 at read length 101 for bulk data and GENCODE v44 at read length 98 for single-cell data. For SCALPEL, index construction consisted of reference generation using cellranger mkref; the prebuilt SCALPEL internal-priming track was excluded.

## Data availability

All processed data generated as part of this study are provided as Supplementary Datasets. ML models are available via Zenodo (DOI: 10.5281/zenodo.12637434). Raw RNA-sequencing data generated in this study are available via GEO (accession numbers GSE271530 and GSE343912).

## Code availability

MPAQT is available at https://github.com/csglab/MPAQT.

## Acknowledgements

We thank Aldo H. Corchado for technical support. This work was supported by the Canadian Institutes of Health Research (CIHR) grant PJT-173317 and resource allocations from Digital Research Alliance of Canada to HSN. MA was supported by a Canada Graduate Scholarships-Master’s award from CIHR. HG is an Era of Hope Scholar (W81XWH-2210121) and supported by grants from the National Cancer Institute (R01CA240984 and R01CA244634). HSN holds a CIHR Canada Research Chair.

## Author contributions

Conceptualization: HSN. Methodology: MA, JW, AS, and HSN. Mathematical derivation: HSN. Code implementation: MA, JW, AS, and HSN. Experiments: BC, AN, and HG. Analysis: MA, JW, AS, LMS, AP, DS, AS, HG, and HSN. Visualization: MA, JW, and HSN. Writing: MA and HSN, with contributions from all authors. Supervision: HG and HSN. Project administration: HSN.

## Competing interests

The authors declare no competing interests.

## Contents of Supplementary Information

### Supplementary Methods

**Supplementary Figure 1.** MPAQT’s performance on (a) salmon and (b) RSEM’s outliers.

**Supplementary Figure 2.** Comparison of quantification tools after widening the range of acceptable Cq values.

**Supplementary Figure 3.** Schematic illustration of the effect of positional biases on the inference of transcript of origin of each read.

**Supplementary Figure 4.** Ground truth vs inferred abundances of endogenous transcripts in the UHRR dataset.

**Supplementary Figure 5.** Characterization of transcripts differentially quantified by MPAQT upon addition of LR data.

**Supplementary Figure 6.** Differentiation of hESCs to neurons. (**a**) Schematic of neuronal differentiation.

**Supplementary Figure 7.** Characteristics of differentially quantified transcripts between SR-only and LR+SR inferences.

**Supplementary Figure 8.** Cell type and disease associations of top genes whose cassette exons are differentially quantified between SR-only and LR+SR analyses in the neuronal differentiation dataset.

**Supplementary Figure 9.** GC and length biases in LR data.

**Supplementary Figure 10.** Example genes showcasing 3’ bias in ONT long reads in the Sequins dataset.

**Supplementary Figure 11.** Isoform- and differentiation stage-specific abundance prediction based on UTR features.

**Supplementary Figure 12.** Pairwise correlation of transcriptomic profiles in ENCODE or LongBench samples.

**Supplementary Figure 13.** Runtime and peak memory requirements for bulk and single-cell transcript quantification and index construction.

**Supplementary Figure 14.** Comparison of inferred exon inclusion and RT-PCR-based measurements in the neuronal differentiation dataset.

**Supplementary Figure 15.** Effect of novel isoforms on transcript quantification.

**Supplementary Figure 16.** Maximum a priori estimation of transcript abundances resolves measurement dropouts.

**Supplementary Figure 17.** Predicting exon inclusion rate from UTR features of associated transcripts.

### Supplementary Data Tables

**Supplementary Data Table 1.** Inference of gene-level expression using SR data from MAQC.

**Supplementary Data Table 2.** Underlying data for evaluation of quantification accuracy in Sequins and UHRR datasets.

**Supplementary Data Table 3.** Gene-level and isoform-level benchmarking metrics for single-cell quantification in CCLE cell line mixtures.

**Supplementary Data Table 4.** RNA-seq and RT-qPCR-based quantifications in hESCs and differentiated cortical neurons.

**Supplementary Data Table 5.** Underlying data for evaluation of quantification accuracy in the LongBench dataset.

**Supplementary Data Table 6.** Data related to ML-based priors for isoform quantification.

## Notes

https://zenodo.org/records/12637435

https://www.ncbi.nlm.nih.gov/geo/query/acc.cgi?acc=GSE271530

https://www.ncbi.nlm.nih.gov/geo/query/acc.cgi?acc=GSE343912

https://github.com/csglab/MPAQT

### Competing Interest Statement

The authors have declared no competing interest.

### Summary of Updates

The methodology has been updated to include the capability for isoform quantification from single-cell data, and to incorporate sequence-to-expression or co-expression based priors. Additional benchmarks are included to show performance across a range of datasets and platforms, and to evaluate computational efficiency.

