## Supplementary Information for "Accurate isoform quantification in bulk and single-cell RNA-seq by integrating multi-platform sequencing data and sequence-to-expression models"

### Supplementary Methods

We use a block-coordinate descent approach to iteratively update individual parameters of the model, including the transcript abundance vector $\boldsymbol{\beta}$, platform-specific library sizes $s_k$, and the effect of confounding biases, represented by vector $\boldsymbol{\gamma}$.

#### Inferring $\boldsymbol{\beta}$

We start by showing how the maximum-likelihood estimate (MLE) of $\boldsymbol{\beta}$ can be obtained in the absence of a log-normal prior on $\boldsymbol{\beta}$. We will then show the inference of the maximum a posteriori (MAP) estimate of $\boldsymbol{\beta}$ with a log-normal prior. As described in the Methods section, in the absence of a prior, for each platform $k$ we have:

$$\forall k \in \{1, \dots, K\} \Big| \boldsymbol{\lambda}_k = \left(\lambda_{k,1}, \dots, \lambda_{k,|U_k|}\right)^{\top} = s_k \boldsymbol{P}_k \boldsymbol{\beta}$$
$$\forall u \in U_k \big| \, n_{k,u} \sim \text{Pois}\left(\lambda_{k,u}\right)$$

Let's define the matrix $\boldsymbol{P}$ as:

$$\boldsymbol{P} = \begin{bmatrix} s_1 \boldsymbol{P}_1 \\ \vdots \\ s_K \boldsymbol{P}_K \end{bmatrix}$$

Then we can rewrite the model as:

$$\boldsymbol{\lambda} = \boldsymbol{P}\boldsymbol{\beta}$$
$$\forall u \in \{U_1, \dots, U_U\} | \, n_u \sim \text{Pois}(\lambda_u)$$

The MLE can be obtained by minimizing the negative log-likelihood function:

$$\widehat{\boldsymbol{\beta}} = \arg\min_{\boldsymbol{\beta}} \sum_{u \in U} (\lambda_u - n_u \log \lambda_u)$$

We use a sequential coordinate-wise descent (SCD) approach to iteratively solve each element $\beta_t$ of $\boldsymbol{\beta}$ ($t \in T$). Consider the current estimate $\beta_t^i$, where $i$ denotes the last iteration of the optimization algorithm. Let's assume that in the next iteration ($i$+1), the updated estimate for $\beta_t$ will be different from the current estimate by $\delta^{[i+1]}$; i.e., the next estimate will be $\beta_t^{[i+1]} = \beta_t^{[i]} + \delta^{[i+1]}$. If the vector of the current predicted OU abundances is $\boldsymbol{\lambda}^{[i]}$, then the next set of predicted fragment abundances is given by:

$$\lambda_u^{[i+1]} = \lambda_u^{[i]} + \delta^{[i+1]} p_{u,t}$$

Therefore:

$$\beta_t^{[i+1]} = \beta_t^{[i]} + \delta^{[i+1]}$$
$$\delta^{[i+1]} = \arg\min_{\delta} \sum_{u \in U} \left[\lambda_u^{[i]} + \delta p_{u,t} - n_u \log\left(\lambda_u^{[i]} + \delta p_{u,t}\right)\right]$$

To solve for $\delta^{[i+1]}$, we take the derivative of the negative log-likelihood function with respect to $\delta$ and set it to zero:

$$\frac{d}{d\delta}\sum_{u\in U}\left[\lambda_u^i+\delta p_{u,t}-n_u\log\left(\lambda_u^i+\delta p_{u,t}\right)\right]=0$$

$$\sum_{u\in U}p_{u,t}-\frac{n_u p_{u,t}}{\lambda_u^i+\delta p_{u,t}}=0$$

To find the root of this function, we use Newton's method to update $\delta$ in each iteration:

$$f(\delta)=\sum_{u\in U}\left[p_{u,t}-\frac{n_u p_{u,t}}{\lambda_u^i+\delta p_{u,t}}\right]=\sum_{u\in U}p_{u,t}\left(1-\frac{n_u}{\lambda_u^i+\delta p_{u,t}}\right)$$

$$\delta^{[i+1]}=\delta^{[i]}-\frac{f\left(\delta^{[i]}\right)}{f'(\delta^{[i]})}$$

$$\delta^{[i+1]}=\delta^{[i]}-\frac{\sum_{u\in U}p_{u,t}\left(1-\frac{n_u}{\lambda_u^i+\delta^{[i]}p_{u,t}}\right)}{\sum_{u\in U}n_u\left(\frac{p_{u,t}}{\lambda_u^i+\delta^{[i]}p_{u,t}}\right)^2}$$

Note that since $\delta^{[i+1]}$ is calculated with respect to the current value of $\beta_t^{[i]}$, then $\delta^{[i]}$ must also be calculated relative to $\beta_t^{[i]}$, which means that $\delta^{[i]}=\beta_t^{[i]}-\beta_t^{[i]}=0$. Therefore:

$$\delta^{[i+1]}=-\frac{\sum_{u\in U}p_{u,t}\left(1-\frac{n_u}{\lambda_u^i}\right)}{\sum_{u\in U}n_u\left(\frac{p_{u,t}}{\lambda_u^i}\right)^2}$$

This provides an iterative procedure where each $\beta_t$ is updated by adding the value of $\delta^{[i+1]}$ from the equation above, followed by updating each $\lambda_u^{[i]}$ (for $u\in U$) by adding $\delta^{[i+1]}p_{u,t}$ to it.

Now, we will modify the equations above to show how the MAP estimate of $\boldsymbol{\beta}$ can be obtained when a log-normal prior is placed on $\boldsymbol{\beta}$:

$$\log\boldsymbol{\beta}\sim\mathcal{N}(\boldsymbol{\theta},\sigma^2\boldsymbol{I})$$

$$\boldsymbol{\lambda}=\boldsymbol{P\beta}$$

$$n_u\sim\text{Pois}(\lambda_u)$$

In this case, the negative log-likelihood function also includes a regularization term that acts to shrink the logarithm of each $\beta_t$ toward the mean of the prior:

$$\widehat{\boldsymbol{\beta}}=\arg\min_{\boldsymbol{\beta}}\left[\frac{1}{2\sigma^2}\sum_{t\in T}(\log\beta_t-\theta_t)^2+\sum_{u\in U}(\lambda_u-n_u\log\lambda_u)\right]$$

Following the same method as above, we can see that for each transcript $t$, its abundance in iteration $i$+1 can be updated as:

$$\beta_t^{[i+1]}=\beta_t^{[i]}+\delta^{[i+1]}$$

$$\delta^{[i+1]} = \arg\min_{\delta}\left[\frac{1}{2\sigma^2}\left[\log\left(\beta_t^{[i]}+\delta\right)-\theta_t\right]^2 + \sum_{u\in U}\left[\lambda_u^{[i]}+\delta p_{u,t} - n_u\log\left(\lambda_u^{[i]}+\delta p_{u,t}\right)\right]\right]$$

Again, using Newton's method and following the same method as above, we can see that:

$$\delta^{[i+1]} = -\frac{\dfrac{\log\beta_t^{[i]}-\theta_t}{\sigma^2\beta_t^{[i]}} + \sum_{u\in U} p_{u,t}\left(1-\dfrac{n_u}{\lambda_u^i}\right)}{\dfrac{-\log\beta_t^{[i]}+\theta_t+1}{\sigma^2\left(\beta_t^{[i]}\right)^2} + \sum_{u\in U} n_u\left(\dfrac{p_{u,t}}{\lambda_u^i}\right)^2}$$

In practice, we have found that in the presence of a log-normal prior, Newton's method occasionally overshoots for some transcripts in some of the early iterations of the optimization algorithm. We detect such overshoot events by examining whether $\delta^{[i+1]}$, as calculated by Newtons's method using the equation above, is outside the range between the value obtained from the MLE estimate and $e^{\theta t}$ (the latter is equivalent to $\log \beta_t^{[i+1]} = \theta_t$, i.e., equal to the mean of the prior). When this occurs, we minimize the negative log-likelihood function using the 'optimize' function in R. This procedure, in practice, resolves the overshoot problem in a few iterations, so that in the subsequent iterations Newton's method can be used without any overshoots.

**The variance of the prior distribution for *β***

We use an adaptive prior, which is iteratively updated based on the distribution of all values $\beta_t$ (for all $t\in T$). In other words, after each iteration *i*, we update the prior variance $\sigma^2$ to be the variance of $\log(\beta)-\gamma$.

**The library size $s_k$**

At each iteration *i*, we update $s_k$ for each platform as:

$$s_k^{[i+1]} = \frac{\sum_{u\in U_k} n_{k,u}}{\sum_{u\in U_k}\sum_{t\in T} p_{k,u,t}\beta_t^{[i]}}$$

**The sources of bias in long-read RNA-seq data (*γ*)**

As described in the Methods section, we model the submatrix $\boldsymbol{P}_k$ matrix for long-read data as:

$$\boldsymbol{P}_k = \mathrm{diag}[\exp(\boldsymbol{C\gamma})]$$

where *k* is the index of the dataset containing long-read counts, and $\boldsymbol{C}$ is a $|T|\times|D|$ matrix representing the value of variables *D*, the potential sources of bias, across $|T|$ transcripts. Therefore, we can model the OU counts observed in dataset *k* as:

$$\lambda_{u\in U_k} = \beta_{t(u)}\sum_{d\in D}\exp\left(c_{u,d}\gamma_d\right)$$

$$n_{u\in U_k} \sim \mathrm{Pois}\left(\lambda_{u\in U_k}\right)$$

Here, $t(u)$ represents the transcript that corresponds to OU *u* (note that there is a one-to-one relationship between OUs in the long-read data and the transcripts). $c_{u,d}$ represents the element $(u,d)$ of matrix $\boldsymbol{C}$. At each iteration *i*, we update $\boldsymbol{\gamma}$ by maximizing the model likelihood using the 'glm' function in R with a Poisson error distribution and log-link, with the long-read counts as the dependent variable, $\boldsymbol{C}$ as the independent variables, and $\boldsymbol{\beta}^{[i]}$ as the offset.

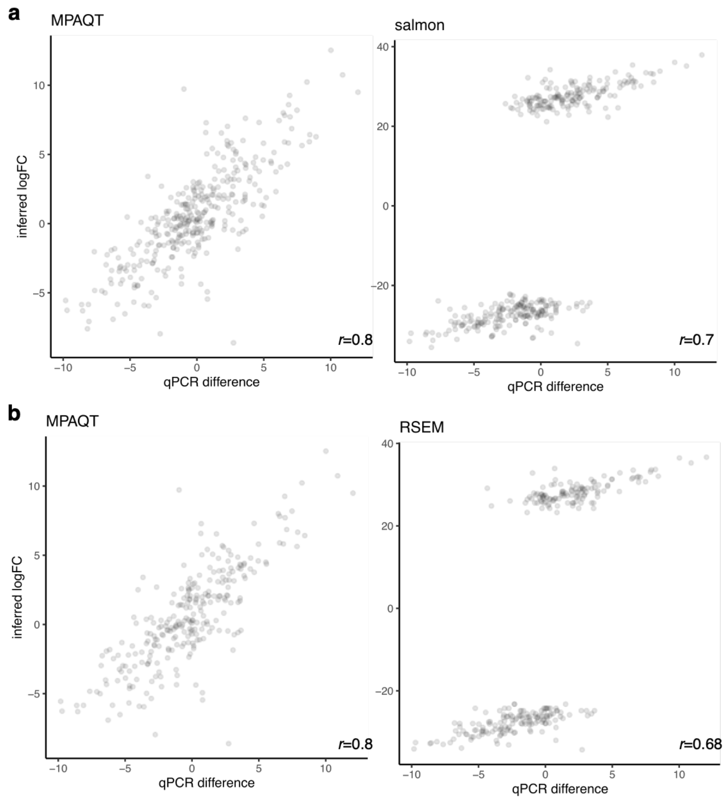


**Supplementary Figure 1.** MPAQT's performance on (a) salmon and (b) RSEM's outliers.

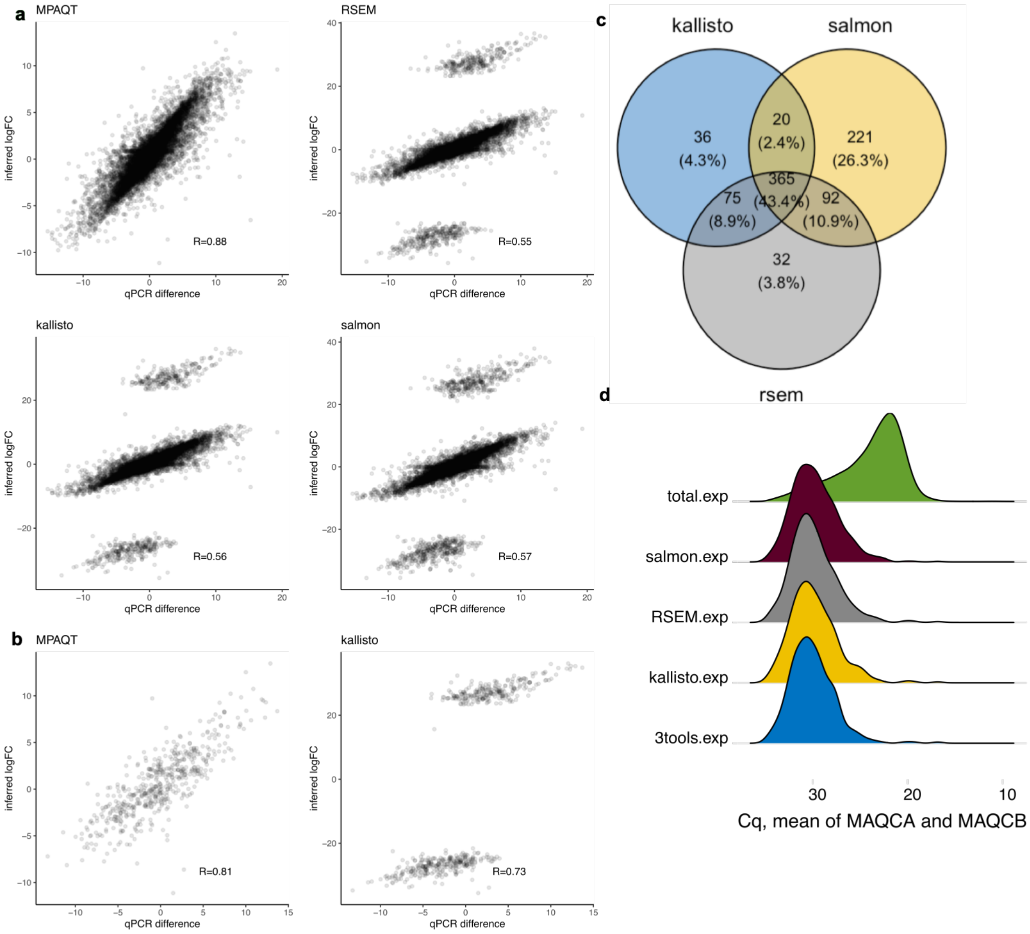


**Supplementary Figure 2.** Comparison of quantification tools after widening the range of acceptable Cq values. Widening the range of Cq to values between 8-35 does not significantly change the MPAQT performance, while the number of outliers for the other three tools almost doubles. The number of genes remaining after filtering increases from 14,956 to 16,104. **(a-d)** Similar to **Figure 1c-f**, respectively, but for the extended Cq range.

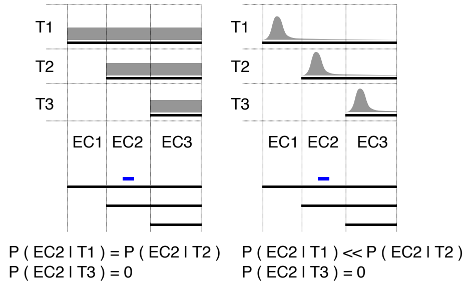


**Supplementary Figure 3.** Schematic illustration of the effect of positional biases on the inference of transcript of origin of each read. Left: a toy example with three isoforms and no positional bias. Right: same toy example, but with a positional bias for transcript termini. In the left example, a read that maps to EC2 has the same probability for isoforms T1 and T2, whereas in the right example, the same read is much more likely to have originated from T2.

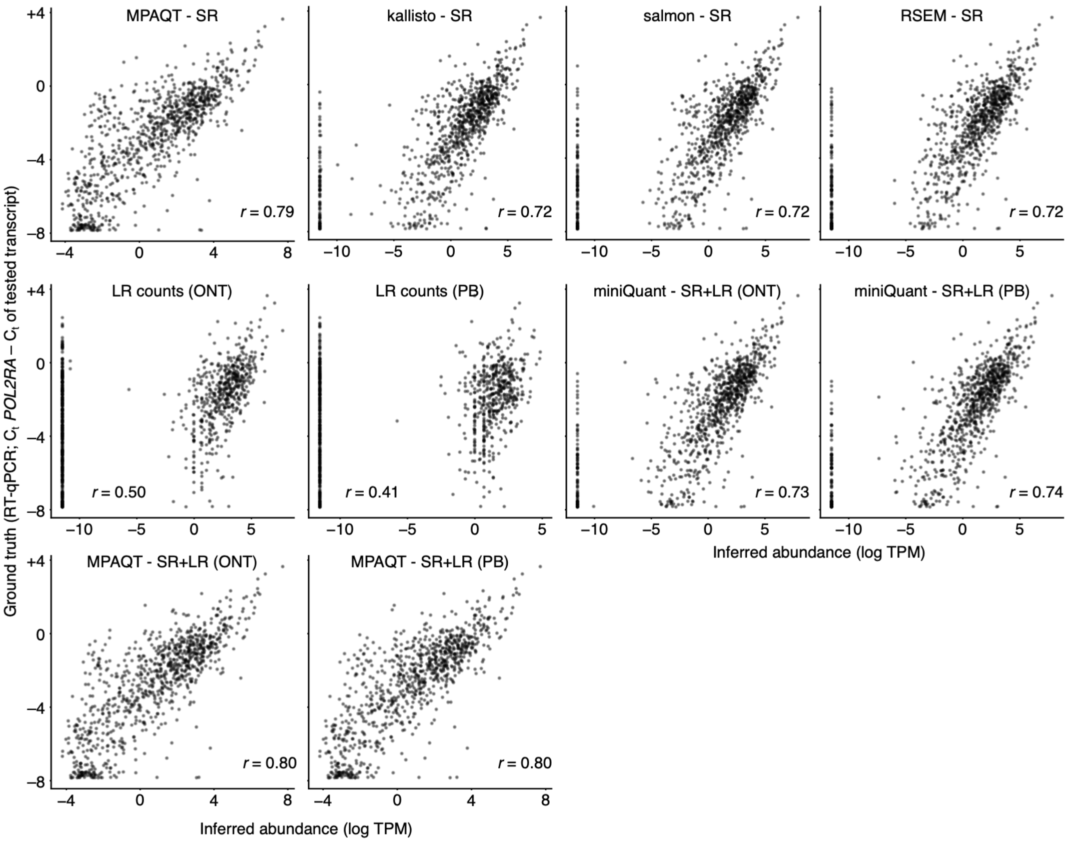


**Supplementary Figure 4.** Ground truth vs inferred abundances of endogenous transcripts in the UHRR dataset. Each inference approach is shown in a separate scatter plot. SR: short-read. LR: long-read. PB: PacBio. ONT: Oxford Nanopore.

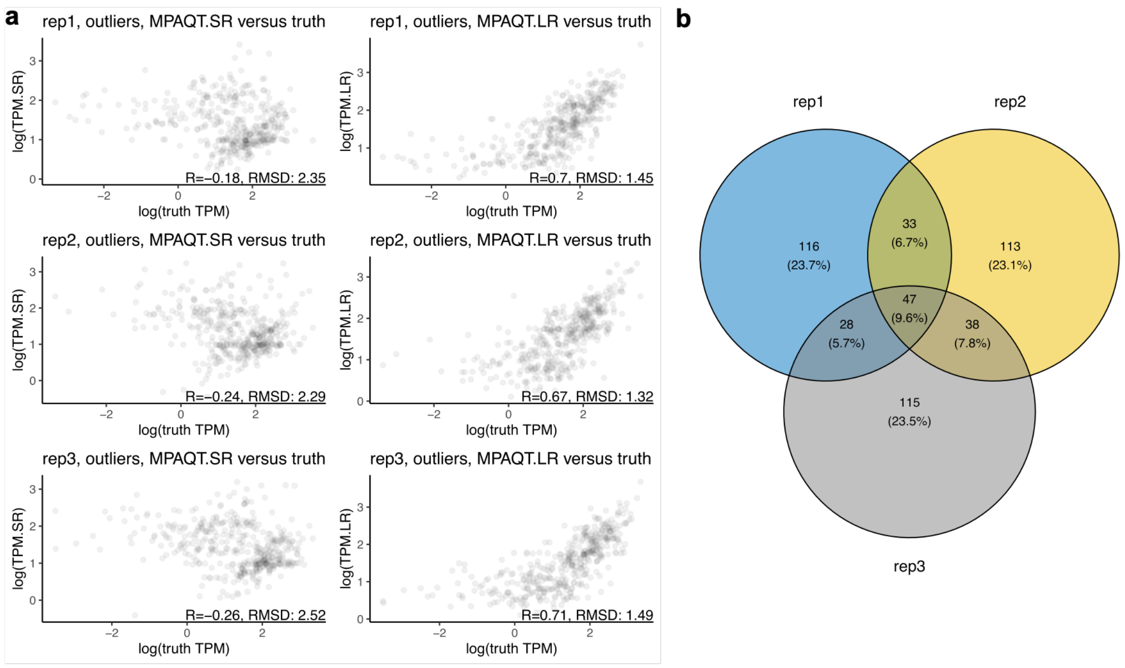


**Supplementary Figure 5.** Characterization of transcripts differentially quantified by MPAQT upon addition of LR data. Transcripts with substantially different inferences were between SR and LR+SR analyses were extracted (outlier analysis based on Mahalanobis distance >6.36, equivalent to upper-tail P<10–10 for normally distributed data), separately for each of the three randomly generated simulated datasets (with exponential distribution of expression). **(a)** For the subset of outlier genes, quantifications correlate better with ground-truth TPMs in MPAQT analysis of LR+SR data than SR alone across all three replicates. **(b)** Venn diagram of the outliers of the three simulated samples, for genes longer than 250bp.

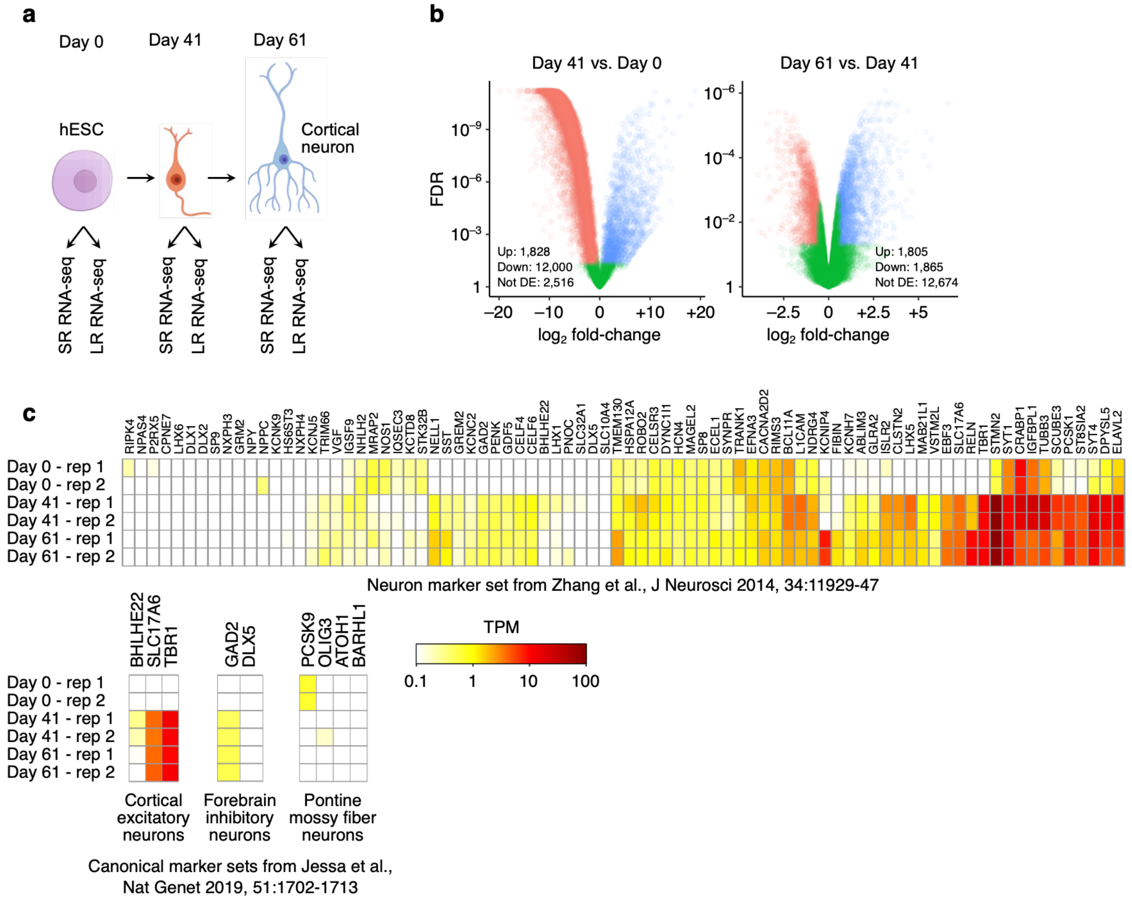


**Supplementary Figure 6.** Differentiation of hESCs to neurons. (**a**) Schematic of neuronal differentiation. Two replicates were collected for each sample, followed by RNA-seq of each replicate using either short-read or long-read sequencing. (**b**) Volcano plots of upregulated (blue), downregulated (red) and non-DE (green) genes between days 0 and 41 (left) and days 41 and 61 (right). (**c**) Heatmaps showing the TPM of neuronal marker genes from ref[1] (top), and different neuronal subclasses from ref[2] (bottom).

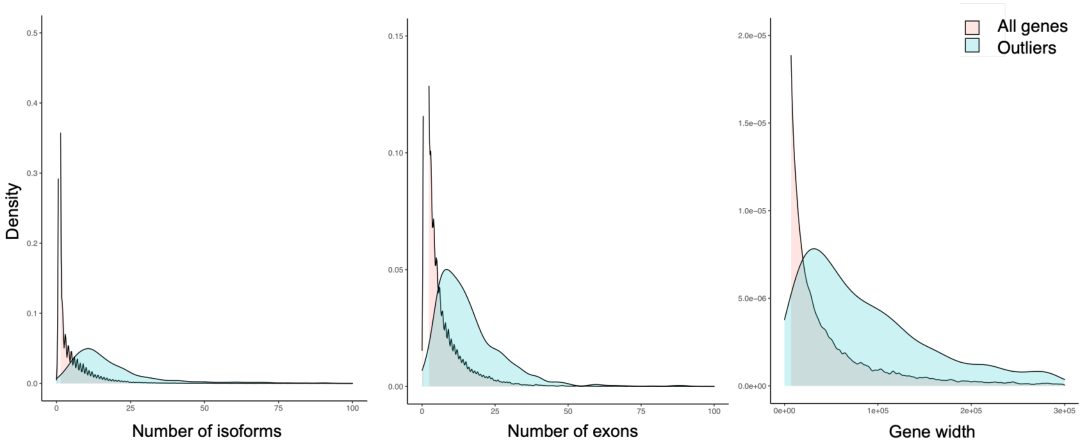


**Supplementary Figure 7.** Characteristics of differentially quantified transcripts between SR-only and LR+SR inferences at day 61 of the neuronal differentiation dataset.

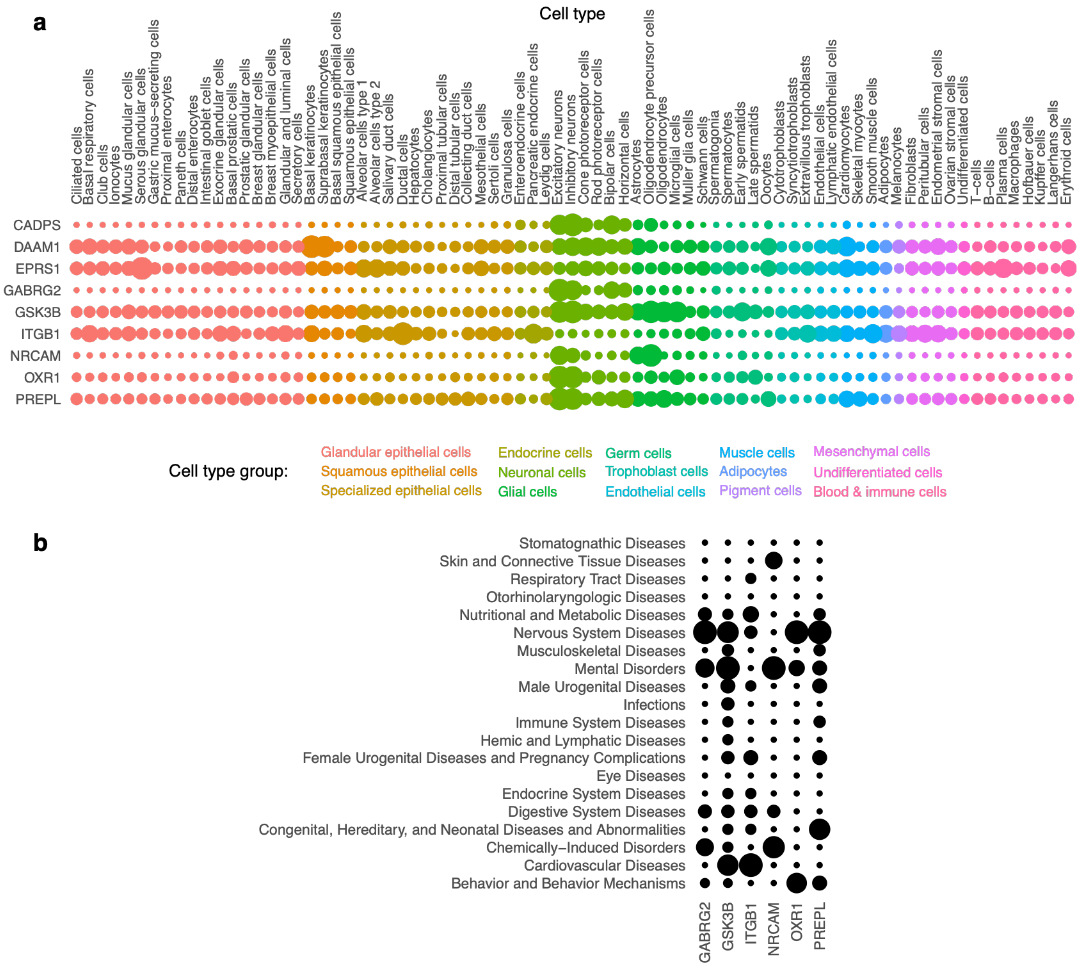


**Supplementary Figure 8.** Cell type and disease associations of top genes whose cassette exons are differentially quantified between SR-only and LR+SR analyses in the neuronal differentiation dataset. (**a**) A dot plot representing the cell type-specific expression, based on RNA single-cell data from Human Protein Atlas[3]. Dot size represents TPM of each gene in each cell type, relative to the cell type with maximum expression of that gene. (**b**) Reported gene-disease associations (GDAs); data from DisGenNet[4]. The rows represent disease classes, and the columns are the genes. The size of each dot represents the number of associations with GDA score >0.1, normalized for each gene to the disease class with the largest number of associations.

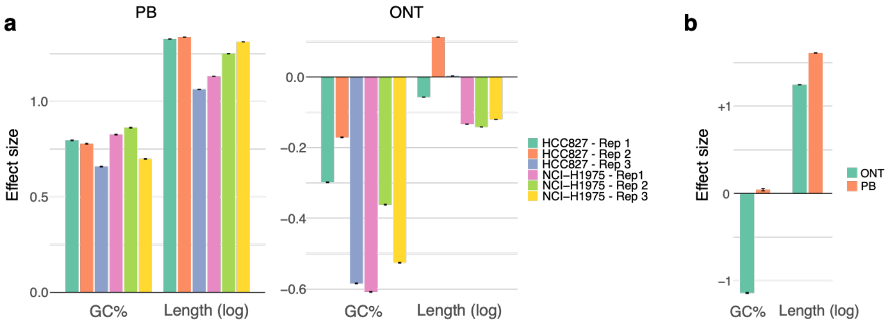


**Supplementary Figure 9.** GC and length biases in LR data. (**a**) The effect of GC content and length on LR counts of endogenous transcripts in the Sequins dataset, estimated by including them as covariates in a Poisson regression with log-scale SR-based inferences as the regressor. PacBio (PB) and Oxford Nanopore (ONT) LR data for each cell line are analyzed and shown separately. (**b**) Same as (a) but for the UHRR dataset.

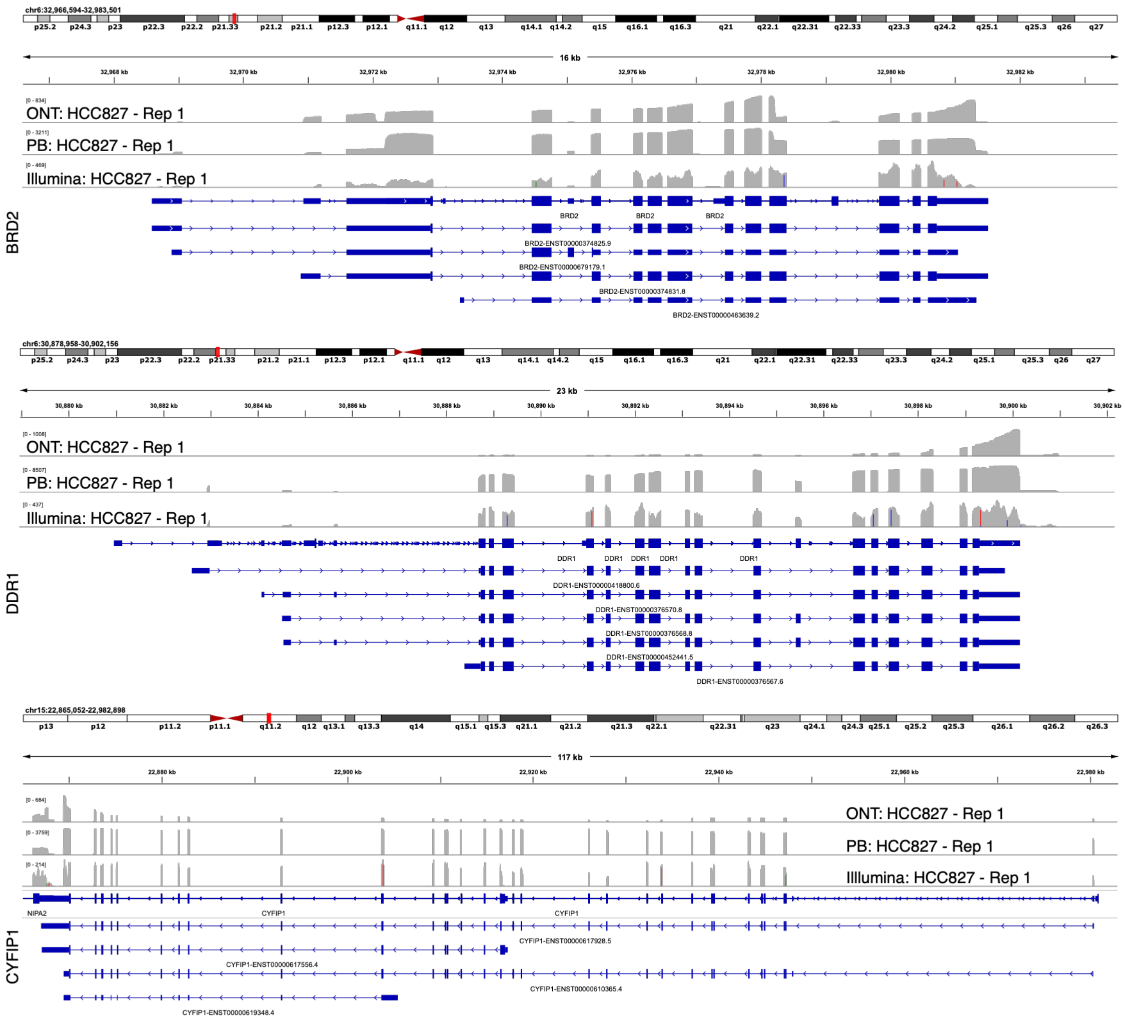


**Supplementary Figure 10.** Example genes showcasing 3' bias in ONT long reads in the Sequins dataset.

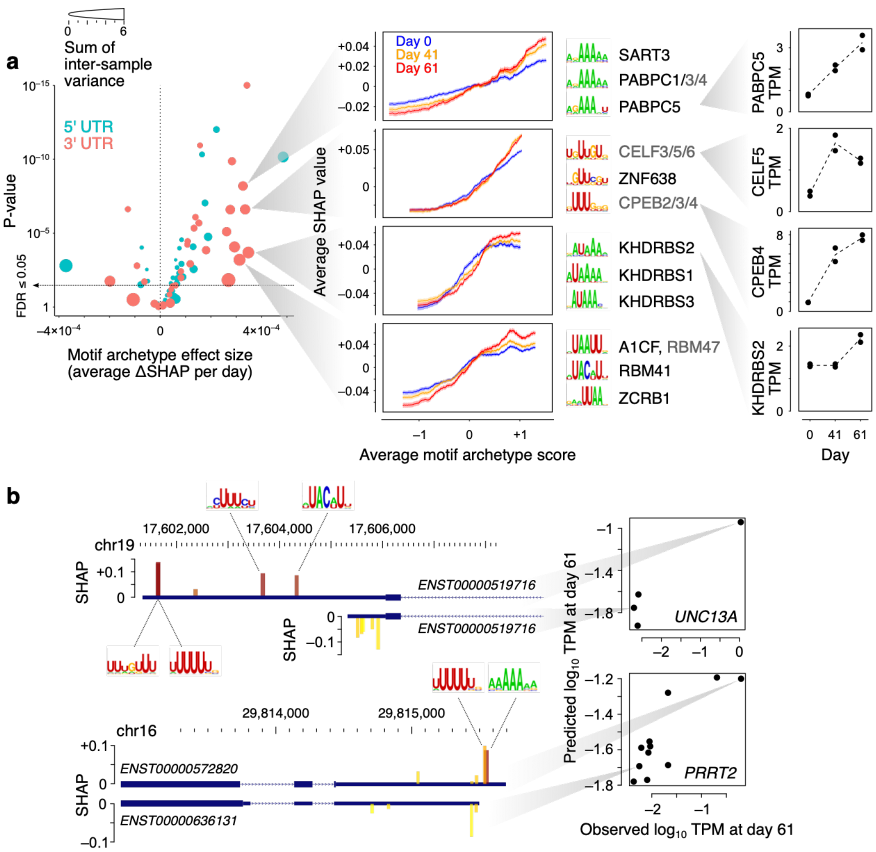


**Supplementary Figure 11.** Isoform- and differentiation stage-specific abundance prediction based on UTR features. (**a**) **Left**: Volcano plot of the differentiation-associated change in SHAP value per motif archetype. For each motif archetype, we tested its contribution to differential mRNA abundances by examining how the relationship between the motif archetype scores and SHAP values change as a function of differentiation time point. For example, for transcripts with high motif archetype scores, if the SHAP value of that motif archetype increases through differentiation, it signifies increased stability of those transcripts due to the contribution of that motif archetype. In order to identify such associations, SHAP values were calculated on held-out transcripts, separately for each time point and each replicate, using the "explain" function from "fastshap" package v0.1.0. Then, for each motif archetype, we concatenated the SHAP values across all transcripts and time point replicates, and fitted a linear regression model of the form $y{\sim}t{+}x{+}t{:}x$, where $y$ is the SHAP value, $t$ is the differentiation time point (0, 41, or 61), and $x$ is the binary variable indicating whether a transcript is among the top 500 transcripts with the largest score for the motif archetype of interest ($x$=1) or not ($x$=0). The coefficient of the interaction term $t{:}x$ represents the change in the SHAP value of top-scoring transcripts as a function of time, which is shown on the x-axis, along with its associated p-value on the y-axis. The size of each circle represents the sum of transcript-wise variances of the SHAP values across time points/replicates. **Middle**: Example motif archetypes with the largest effect sizes and sample-to-sample variances. For each motif archetype, the moving average chart of SHAP vs. motif archetype score is shown (transcripts were sorted by their motif archetype scores, following by mean calculation over sliding windows of 500 transcripts). Each curve represents one time point. The shaded areas correspond to the standard error of mean of SHAP values per sliding window. The top three motifs associated with each motif archetype are shown next to each chart, along with the RBPs that recognize each motif (RBPs shown in grey are inferred to recognize the motif based on homology[19]). **Right**: Gene-level TPM profiles for example RBPs across differentiation time points. (**b**) Example isoforms of *UNC13A* (top) and *PRRT2* (bottom). For each gene, one dominant isoform and one low-abundance isoform is shown, along with the top five motifs whose presence in the 3' UTR explains the higher abundance of the dominant isoform. For each motif, the position of the best-matching sequence in each isoform is shown, along with the SHAP value of the associated motif archetype (shown with the bar height) and the motif hit score (yellow: low-scoring hit; red: high-scoring hit). The predicted and observed abundances of the highlighted isoforms (along with other isoforms of each gene) are shown in the scatterplot on the right.

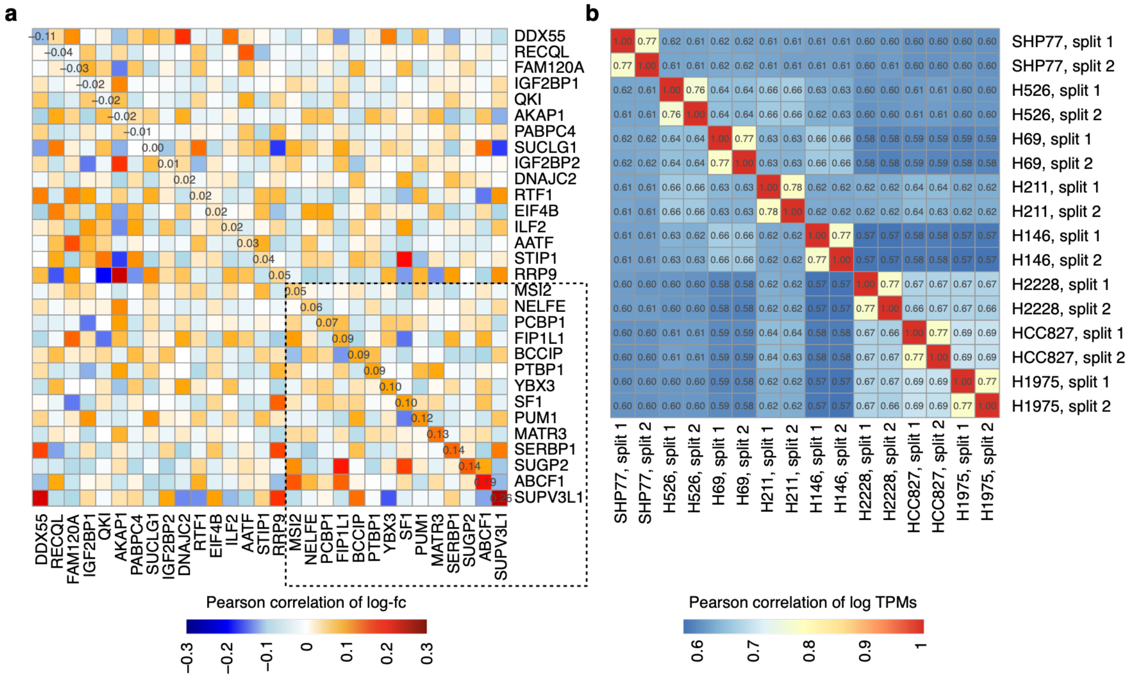

**Supplementary Figure 12.** Pairwise correlation of transcriptomic profiles in ENCODE or LongBench samples. (**a**) Correlation of transcriptomic response to RBP perturbations in K562 (rows) and HepG2 (columns). In each pairwise comparison, Pearson correlation is calculated using mean log fold-change between biological replicates relative to the mean of all perturbations (across transcripts with a TPM of 1 in at least 5% of all samples). The dotted square highlights the RBPs whose transcriptomic responses have correlation >0.05 between the two cell lines. (**b**) Correlation between in silico technical replicates of bulk RNA-seq data from the LongBench dataset. To partition the bulk short-read data into simulated technical replicates, we used the "partition.sh" tool from bbmap (version 39.06) with setting ways=2 to split each paired-end FASTQ into 2 replicates with equal numbers of reads, which were then analyzed by kallisto to quantify isoform abundances. The heatmap shows the pairwise correlations across protein-coding transcripts.

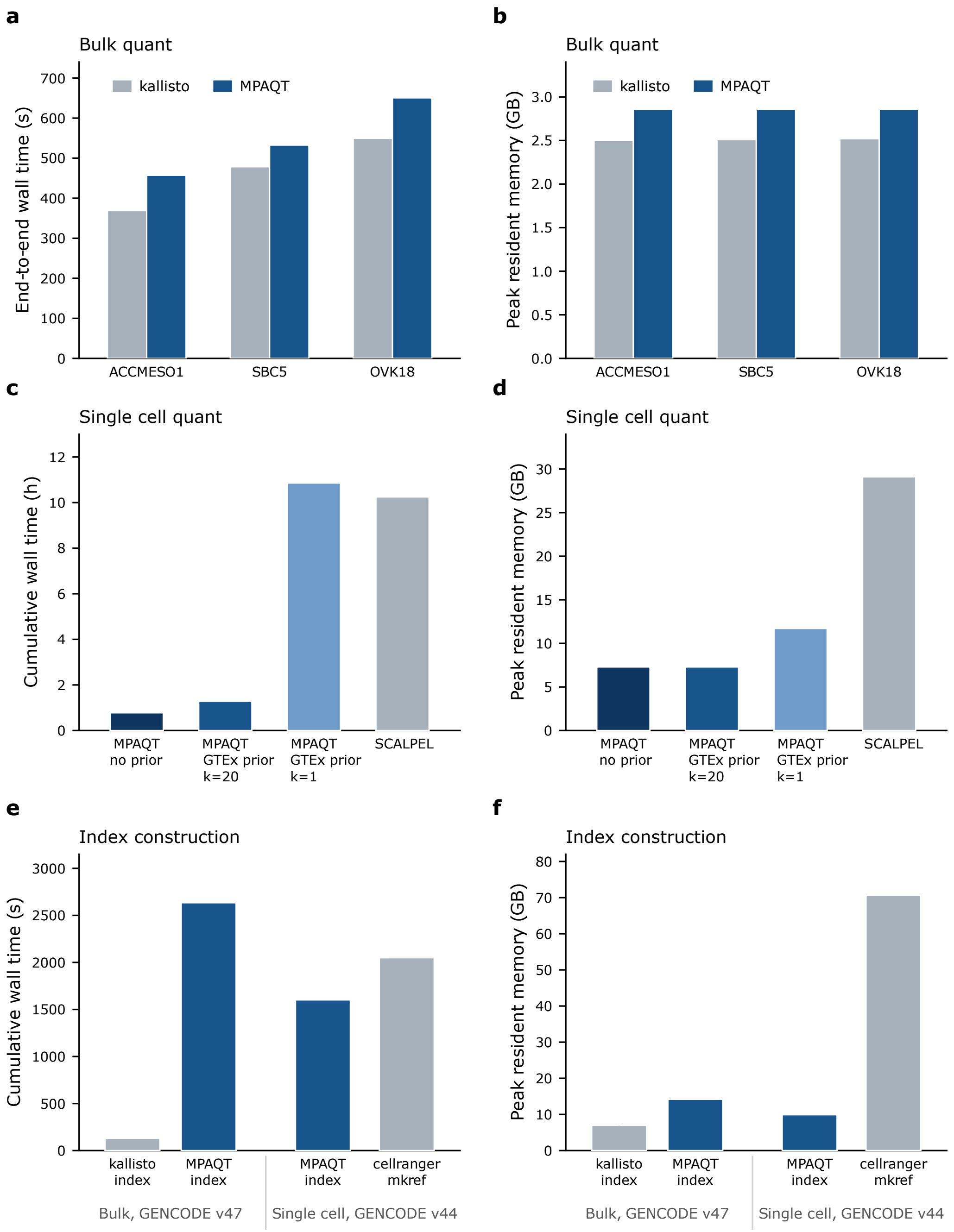


**Supplementary Figure 13.** Runtime and peak memory requirements for bulk and single-cell transcript quantification and index construction. (**a-b**) runtime (a) and peak memory (b) for MPAQT and kallisto across three CCLE bulk RNA-seq libraries: SRR8615406 (ACCMESO1), SRR8616102 (SBC5), and SRR8615880 (OVK18). (**c-d**) runtime (c) and peak memory (d) for MPAQT and SCALPEL across 24 cell-line clusters. MPAQT was run without sequence-informed LR prior and with a GTEx prior refitted every iteration (k=1) or every 20 iterations (k=20). (**e-f**) runtime (e) and peak memory (f) for index construction, measured separately and excluded from quantification runtimes.

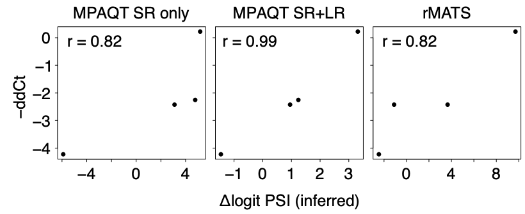


**Supplementary Figure 14.** Comparison of inferred exon inclusion and RT-PCR-based measurements in the neuronal differentiation dataset. To run rMATS, adaptor trimming was performed using Trimmomatic v0.35 , followed by alignment with STAR v2.5.0c (genome version GRCh38, Ensembl87.sjdbOverhang99). rMATS analysis was performed with rmats v4.3.0, with parameters -t paired, --readLength 75, and --variable-read-length, using GTF reference GENCODE v37. For all tools, Δlogit-PSI was calculated as $\Delta\text{logit}(\Psi) = \text{logit}(\Psi\_\text{Day61}) - \text{logit}(\Psi\_\text{Day0})$, where $\text{logit}(\Psi) = \ln[(\Psi + \varepsilon) / (1 - \Psi + \varepsilon)]$, $\varepsilon = 0.0001$. Only four exons with RT-PCR measurements were also present in rMATS output, which are included in this plot.

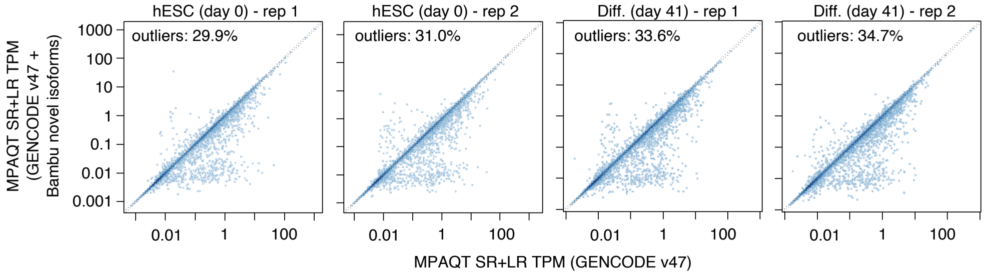


**Supplementary Figure 15.** Comparison of isoform abundances inferred based on GENCODE v47 or based on GENCODE v47 augmented with novel isoforms identified by Bambu from neuronal differentiation LR data. Only GENCODE v47-annotated isoforms are shown in these plots, limiting to those from genes that contained novel isoforms. The percentage of isoforms with >10% change in abundance as a result of inclusion of novel isoforms is shown on each plot.

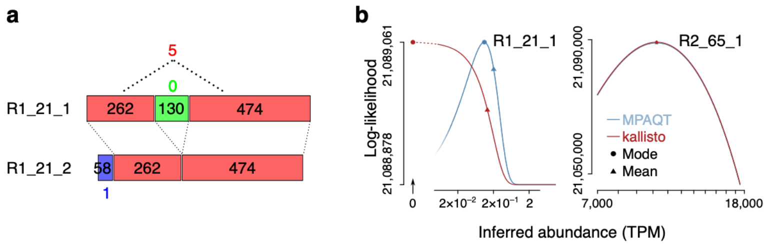


**Supplementary Figure 16.** Maximum a priori estimation of transcript abundances resolves measurement dropouts. (**a**) Two example Sequin isoforms and their associated read counts (in HCC827 experiment). Each unique color represents one EC. Black numbers represent exon lengths, while colored numbers represent observed counts for color-matched ECs. (**b**) Left: the likelihood profile for R1_21_1 based on kallisto's model (red) or MPAQT's model (cyan). Circles and triangles represent mode and mean of each distribution, respectively. Right: the likelihood profiles for R2_65_1, an example Sequin transcript that has non-zero reported TPM in kallisto analysis (the two profiles overlap almost perfectly, hence appear as one).

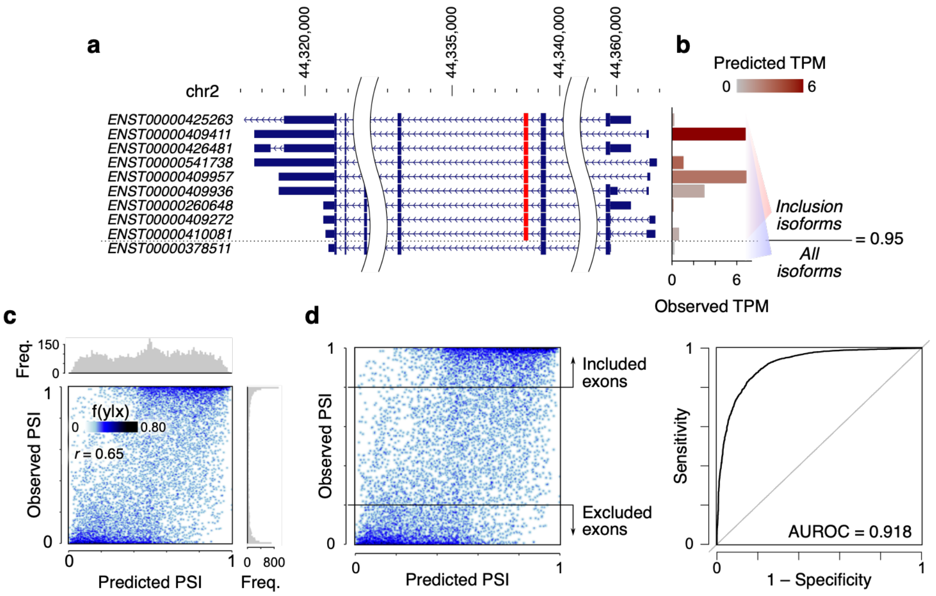


**Supplementary Figure 17.** Predicting exon inclusion rate from UTR features of associated transcripts. (**a**) The isoforms associated with inclusion or exclusion of an example cassette exon (shown in red) for gene *PREPL*. (**b**) The observed TPM of each isoform is shown using the bar graph, with the color gradient specifying the predicted TPM. (**c**) The scatter plot of predicted vs. observed PSI. Each point represents one cassette exon in one time point/replicate (cassette exons for which the sum of TPMs of inclusion and exclusion isoforms was <1 were excluded). (**d**) Same scatterplot as (c) is shown on the left, with horizontal lines representing exons with observed PSI >0.8 ("positive" observations) or PSI <0.2 ("negative" observations). These positive and negative observations are used to construct the classification ROC curve shown on the right.
